# Identification of Aryl Hydrocarbon Receptor as a Barrier to HIV-1 Infection and Outgrowth in CD4^+^ T-Cells

**DOI:** 10.1101/2022.10.17.512596

**Authors:** Debashree Chatterjee, Yuwei Zhang, Tomas Raul Wiche Salinas, Christ-Dominique Ngassaki-Yoka, Huicheng Chen, Yasmine Smail, Jean-Philippe Goulet, Brendan Bell, Jean-Pierre Routy, Petronela Ancuta

## Abstract

The Aryl hydrocarbon receptor (AhR) identifies “*non-pathogenic*” Th17-polarized CD4^+^ T-cells in autoimmune models. Thus, we explored whether AhR restricts HIV-1 in Th17-cells, consistent with its antiviral role in macrophages. AhR-specific CRISPR/Cas9-mediated knockout and pharmacological blockade decreased AhR target gene expression (CYP1A1/IL-22/IL-17A/IL-10/ ITGB7), while increasing HIV-1 replication in CD4^+^ T-cells. Pharmacological AhR activation caused opposite effects. AhR agonism/antagonism modulated HIV-1 replication mainly in Th17/Th22-polarized CCR6^+^CD4^+^ T-cells. Single-round VSV-G-pseudotyped HIV-1 infection demonstrated that AhR acts at post-entry levels, with AhR blockade increasing the efficacy of early/late reverse transcription steps and subsequently integration/translation. In viral outgrowth assay, the AhR blockade boosted the detection of replication-competent viral reservoirs in CD4^+^ T-cells of people living with HIV-1 (PLWH) receiving antiretroviral therapy (ART). Finally, RNA-Sequencing revealed genes/pathways modulated by AhR blockade in CD4^+^ T-cells of ART-treated PLWH, with known HIV-1 interactor activities (NCBI HIV Interactor Database) and AhR responsive elements in their promoters (ENCODE). Among them, HIC1, a repressor of Tat-mediated HIV-1 transcription and a tissue-residency inducer, represents a putative AhR mechanism of action. These results demonstrate that AhR governs an antiviral transcriptional program in CD4^+^ T-cells and point to the use of AhR inhibitors to boost viral outgrowth in “*shock and kill*” HIV-1 remission/cure strategies.

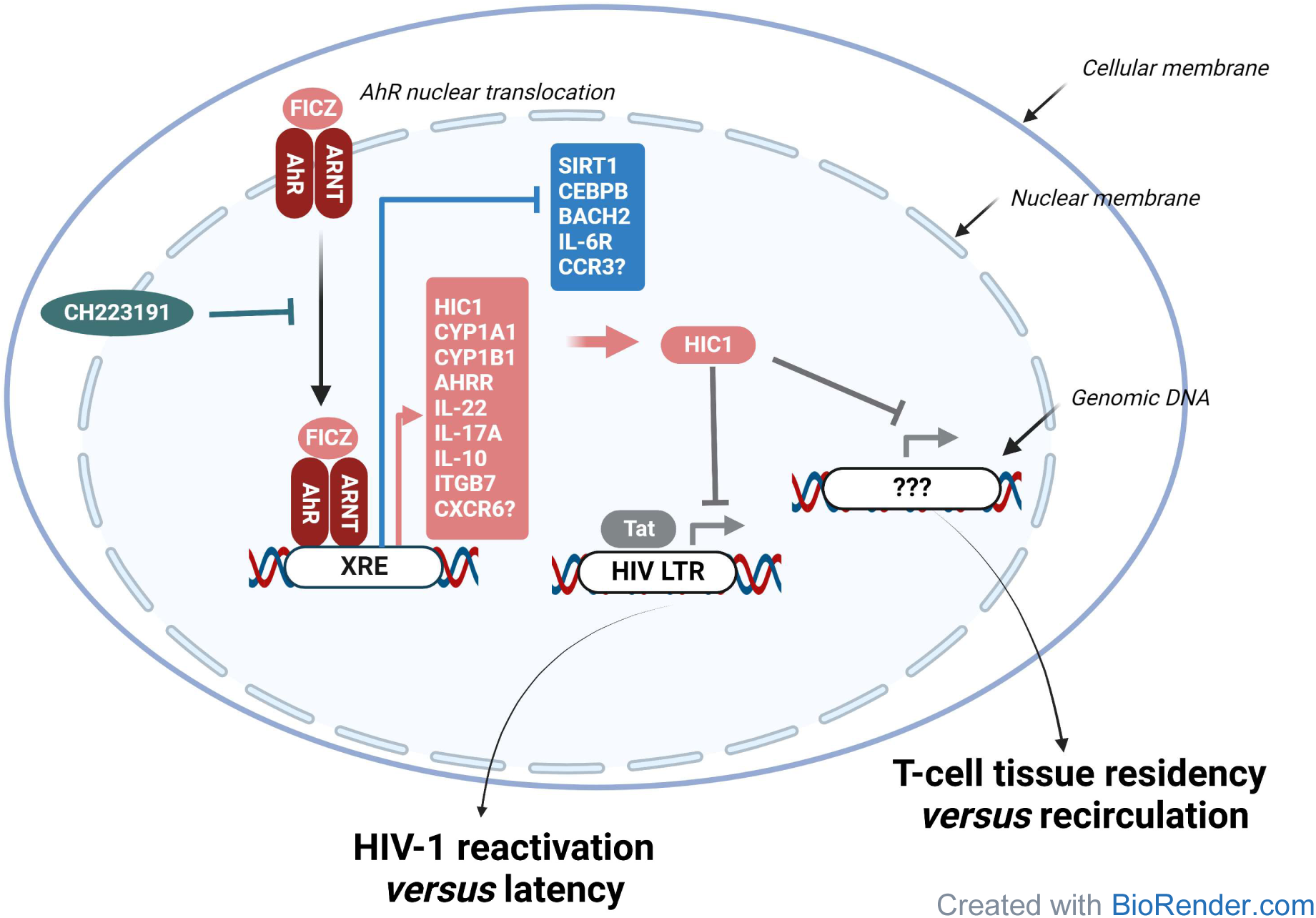
Model of AhR-mediated transcriptional reprogramming with implications for “*silent*” HIV-1 reservoir persistence and gut homing/residency. RNA-Sequencing revealed genes sets modulated by AhR blockade in CD4^+^ T-cells of ART-treated PLWH, with known HIV-1 interactor activities (NCBI HIV Interactor Database) and AhR responsive elements in their promoters (ENCODE). Among them, HIC1, a repressor of Tat-mediated HIV-1 transcription and a tissue-residency regulator, represents a putative AhR mechanism of action. These results support a model in which AhR activation favors the gut homing and residency *via* the induction of ITGB7 and CXCR6 expression, respectively, and fuels the persistence of ‘*silent*” HIV-1 reservoirs in CD4^+^ T-cells of ART-treated PLWH. At the opposite, pharmacological AhR blockade facilitates viral outgrowth, and by interfering with tissue residency, likely promotes the mobilization of « reactivated » reservoir cells from deep tissues into the circulations.

**BRIEF SUMMARY:** We identified the aryl hydrocarbon receptor as a barrier to HIV-1 infection/outgrowth in Th17-polarized CD4^+^ T-cells and a novel therapeutic target in HIV-1 cure/remission interventions.

## INTRODUCTION

Anti-retroviral therapy (ART) transformed HIV-1 infection into a manageable chronic disease and increased the life expectancy of people living with HIV-1 (PLWH). However, ART does not cure HIV-1 and immune competence is not completely restored (1, 2). Major barriers to HIV-1 remission/cure during ART include the persistence of viral reservoirs (VR) in long-lived CD4^+^ T-cells (3, 4), residual HIV-1 transcription in VR cells (5–7), and the non-restoration of intestinal barrier functions (8, 9). Subsequently, ART-treated PLWH experience chronic immune activation, immune metabolism deregulation, and increased risk for non-AIDS comorbidities, such as cardiovascular disease (10, 11). In this context, new therapeutic strategies are required for HIV-1 remission/cure.

The intestinal barrier alterations in PLWH coincide with the massive depletion of CD4^+^ T-cells from gut-associated lymphoid tissues (GALT), a depletion not restored by ART (8, 9). Despite their paucity, GALT-infiltrated/resident CD4^+^ T-cells represent a major pool of cells carrying HIV-1 reservoirs during ART (12). Among CD4^+^ T-cell subsets that play an important role in intestinal homeostasis, cells producing the hallmark cytokines IL-17A/F (Th17) and IL-22 (Th22) are particularly targeted by HIV-1 for infection and subsequently depleted (8, 9, 13). The preferential loss of Th17/Th22 cells is also observed during pathogenic SIV infection, but not in non-pathogenic SIV infection models and in PLWH with virological control in the absence of ART, where Th17/Th22 cells are preserved at mucosal sites (14–18). This emphasizes the importance of Th17/Th22 cells in limiting HIV-1 disease progression. Therefore, the design of novel interventions designed to protect Th17/Th22 cells from infection/depletion represents a priority and requires a detailed understanding of molecular mechanism regulating HIV-1 permissiveness in these cells.

The differentiation and effector functionality of Th17 cells are governed by the master regulator transcription factor (TF) *retinoic acid orphan receptor gamma t* (RORγt in mice; RORC2 in humans) (19, 20). RORγt/RORC2 represents a target for drugs designed to treat autoimmune conditions in which Th17 cells exert deleterious functions (21–23). Of particular notice, recent studies by our group identified RORC2 as a positive regulator of HIV-1 replication in Th17 cells (24). However, only a fraction of Th17 cells is permissive to integrative HIV-1 infection (8, 9, 13), indicative that Th17 cells resistant to HIV-1 infection exist. It is noteworthy that, in the context of autoimmunity, two types of Th17 cells were identified, “*pathogenic*” and “*non-pathogenic*”, based on their ability to promote experimental autoimmune encephalitis (EAE) upon adoptive transfer (25). Recent studies demonstrated the intestinal origin of pathogenic Th17 cells causing EAE (26), thus supporting the crosstalk between gut and distal tissues (*e.g.,* brain). Also, “*non-pathogenic*” Th17 cells were identified in both mice and humans, and documented to express a unique molecular signature including the *aryl hydrocarbon receptor* (AhR) (25, 27). The role of AhR in restricting HIV-1 permissiveness in Th17 cells, thus rendering them “*non-pathogenic*”, remains unknown.

The AhR is a ligand-dependent nuclear receptor, initially identified as the dioxin receptor (28, 29). It is a member of the basic-helix–loop–helix (bHLH)/ Per–Arnt–Sim (PAS) family of proteins that interacts with a broad range of xenobiotic or natural ligands such as tryptophan metabolites, dietary products, and microbiota-derived factors (30–32). In the absence of ligands, AhR resides in the cytoplasm in an inactive state due to its association with different chaperons (*i.e.,* HSP90, XAP2, p23) (32). Upon ligand binding, AhR undergoes a conformational change that allows the interaction with the AhR nuclear translocator (ARNT) and the formation of the AhR/ARNT heterodimer. The AhR/ARNT heterodimer subsequently translocates into the nucleus and binds onto xenobiotic responsive elements (XRE) on the promoter of specific genes, thus regulating their transcription (32). Among AhR regulated genes, the xenobiotic-metabolizing enzyme CYP1A1 (cytochrome P450 family 1 A1), AHRR (AhR repressor), and TIPARP [2,3,7,8-tetrachlorodibenzo-p-dioxin poly(ADP-ribose) polymerase] are involved in the negative feedback of AhR signaling *via* AhR ligand degradation, AhR transcriptional repression, and AhR degradation by ribosylation, respectively (32, 33). Thus, AhR is an important nuclear receptor that adapts transcriptional profiles of cells mainly present at barrier surfaces to changes in the environment (30–32).

Pioneering studies demonstrated that AhR is specifically expressed by Th17-polarized CD4^+^ T-cells from mice and humans (34), with AhR activation promoting effector functions (35–37), such as the production of IL-22 in Th22 cells, as well as IL-10 in regulatory T-cells (Tregs) (34, 35, 38-40). Of particular notice, AhR mediates the trans-differentiation of Th17 cells into Tregs during the resolution of inflammation, a process dependent on AhR-mediated IL-10 production (41). Moreover, AhR was reported to transcriptionally regulate the expression of the gut-homing integrin beta 7 (ITGB7) in macrophages (42), indicative of an AhR role in regulating gut tropism/residency. Furthermore, a large body of evidence identified AhR-expressing CD4^+^ T-cells act as important gatekeepers, mainly by producing lineage-specific cytokines (*e.g.,* IL-17A/F, IL-22) that act on epithelial cells to strengthen their barrier functions (32). Finally, AhR activation was reported to block HIV-1 replication in macrophages (43) thus, raising the possibility that AhR restricts HIV-1 infection in CD4^+^ T-cells as well, especially in Th17 cells, which represent major HIV-1 infection targets (8, 9, 13).

Herein, we explored the role of AhR in modulating HIV-1 replication/outgrowth in primary CD4^+^ T-cells isolated from the peripheral blood of ART-treated PLWH and uninfected participants. Our results provide evidence supporting a model in which AhR governs an antiviral program that, in the context of specific AhR ligands derived from microbiota, diet and tryptophane catabolism (30–32), may reduce the susceptibility to HIV-1 acquisition during primary infection. Once the infection is established, the activation of the AhR pathway may limit HIV-1 outgrowth from VR cells of ART-treated PLWH, therefore preventing VR sensing and depletion by the immune system. Finally, our results support the use of pharmacological AhR inhibitors, a new class of drugs currently tested in clinically for cancer (44, 45), to boost VR reactivation in “*shock and kill*” HIV-1 remission/cure strategies.

## RESULTS

### CRISPR/Cas9-mediated AhR knockout reduces IL-17A, IL-22 and IL-10 production and facilitates HIV-1 replication *in vitro*

To explore the role of AhR in modulating HIV-1 permissiveness in CD4^+^ T-cells, we first explored AhR expression/activation upon TCR triggering. Maximal AhR mRNA and protein expression in memory CD4^+^ T-cells was observed at day 1 and day 3 post-TCR triggering, respectively (<u>Supplemental Figure 1A-B</u>). Interestingly, AhR expression was not induced by PHA activation (<u>Supplemental Figure 1C</u>), likely explaining findings by Kueck *et al.* that HIV-1 replication in PHA-activated CD4^+^ T-cells was AhR-independent (43). Of note, AhR expression in TCR-activated CD4^+^ T-cells coincided with the expression of CyP1A1 mRNA, a AhR-specific target gene and a component of the cytochrome p450 family (32), that culminated at day 3 post-TCR triggering (<u>Supplemental Figure 1D</u>). Thus, TCR triggering in CD4^+^ T-cells leads to AhR expression/activation in memory CD4^+^ T-cells, consistent with the presence of AhR ligands in culture media, as previously reported (37).

To determine the effect of TCR-mediated AhR expression/activation on HIV-1 replication in CD4^+^ T-cells, the CRISPR/Cas9 method for AhR knockout (KO) was optimized in primary CD4^+^ T-cells (Figure 1A), using a published protocol (46). TCR-activated memory CD4^+^ T-cells electroporated using AhR specific (AhR crRNA) compared to nonspecific (negative crRNA) guide RNA resulted in an efficient AhR silencing, as demonstrated by western blotting visualization of AhR protein expression (Figure 1B). The gene editing activity of AhR guide RNA was confirmed by T7 endonuclease assay (Figure 1C). At day 4 post-electroporation, the CRISPR/Cas9-mediated AhR KO resulted in decreased expression of the gut-homing molecule integrin beta 7 (ITGB7) (<u>Supplemental Figure 2A-B</u>), consistent with results in macrophages (42). The electroporation with AhR crRNA and negative crRNA did not affect cell viability (<u>Supplemental Figure 2C</u>). Together, these results demonstrate that CRISPR/Cas9-mediated AhR KO was successful in primary memory CD4^+^ T-cells.

**Figure 1:**
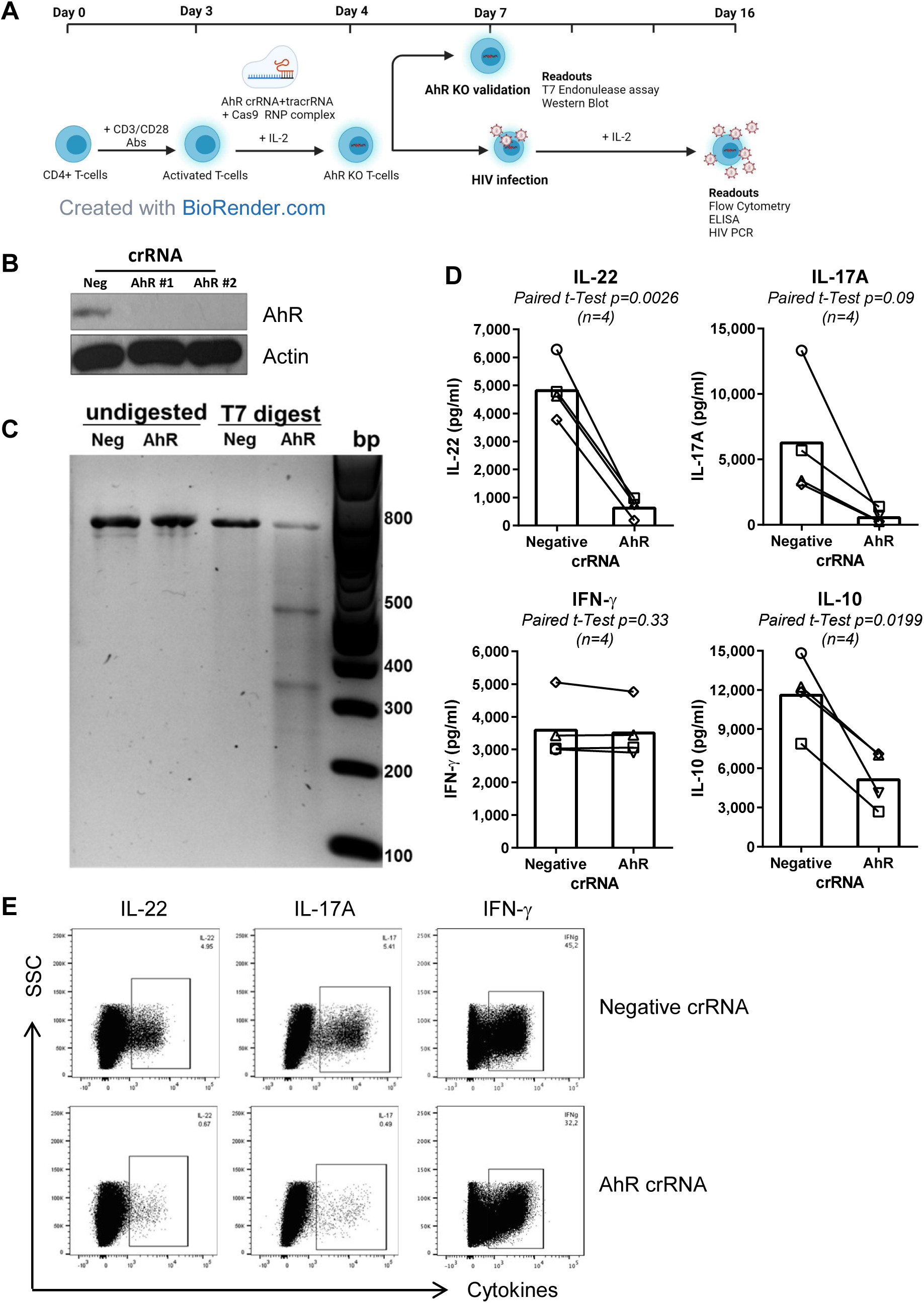
CRISPR/Cas9-mediated AhR KO reduces IL-22, IL-17A and IL-10 production in primary CD4^+^ T-cells. **(A)** Shown is the experimental flow chart of CRISPR/Cas9 ribonucleoprotein (RNP)-mediated AhR KO performed in primary CD4^+^ T-cells. Briefly, memory CD4^+^ T-cells isolated from PBMCs of HIV-uninfected individuals by negative selection using magnetic beads were stimulated with CD3/CD28 Abs for 3 days. Cells were then electroporated with the AhR-targeting crRNA (AhR crRNA) or the control CRISPR (negative crRNA) in the presence of the trans-activating CRISPR RNA (tracrRNA). **(B-C)** Four days after electroporation, cells were analyzed by western blotting **(B)** and the T7 endonuclease assay **(C)** to determine the efficacy of AhR KO at protein and DNA levels, respectively. **(D-E)** In parallel, one day after electroporation, cells were exposed to HIV_THRO_ for 3 hours, the unbound virus was removed by extensive washing and cells were cultured for 9 days in the presence of recombinant human IL-2 (5 ng/ml). At day 9 post-infection, supernatants were collected, and levels of IL-22, IL-17A, IFN-γ, and IL-10 were quantified by ELISA **(D)**. In parallel, cells were stimulated with PMA (50 ng/ml) and Ionomycin (1 µg/ml)for 2 hrs at 37 °C followed by the addition of Brefeldin A (2 µg/ml) and cells are incubate another 4hours at 37 °C and the intracellular expression of IL-22, IL-17A, and IFN-γ was measured by flow cytometry **(E)**. Experiments in **B-C** and **D-E** were performed with cells from n=2 and n=4 HIV-uninfected individuals, respectively. Paired t-Test p-values are indicated on the graphs in panel **E**.

The CRISPR/Cas9-mediated AhR KO protocol (Figure 1A) was further used to explore the AhR-dependent modulation of HIV-1 permissiveness in primary CD4^+^ T-cells. For this, memory CD4^+^ T-cells from HIV-uninfected individuals were electroporated with AhR-crRNA and negative-crRNA and exposed one day later to the transmitted/founder HIV-1 strain (HIV_THRO_). Cells were further cultured in the presence of IL-2 for 9 days to monitor cytokine production and HIV-1 replication. Considering the evidence that AhR transcriptionally regulates the expression of IL-17A and IL-22 (34, 35, 37, 38), as well as IL-10 expression (39), the production of IL-22, IL-17A and IL-10, but not IFN-γ, was strongly downregulated in AhR-crRNA compared to negative-crRNA electroporated CD4^+^ T-cells, as measured by ELISA at day 9 post-infection (Figure 1D). Also, upon PMA/Ionomycin stimulation, AhR-crRNA compared to negative-crRNA electroporated CD4^+^ T-cells expressed lower levels of IL-22 and IL-17A, but similar levels of IFN-γ, as measured by ICS and flow cytometry (Figure 1E). These results are indicative of an efficient AhR functional interference at day 9 post-infection.

Of particular importance, CRISPR/Cas9-mediated AhR KO significantly increased replication of the transmitted/founder HIV-1 strain THRO (HIV_THRO_) (Figure 2), an HIV-1 molecular clone documented to exhibit peculiar infectivity and resistance to type I IFN-mediated antiviral immunity (47). This effect was observed by measuring levels of HIV-DNA integration at day 3 post-infection (Figure 2A), as well as HIV-p24 levels in cell-culture supernatants at days 3, 6 and 9 post-infection (Figure 2B-C). A similar increase in HIV-DNA integration was observed when AhR silencing was induced in TCR-activated CD4^+^ T-cells with AhR (siAhR) *versus* non-targeting small interfering RNA (siNT) and cells were exposed to the replication-competent CCR5-tropic HIV_NL4.3BaL_ strain (<u>Supplemental Figure 3</u>). These results demonstrate that AhR acts as a negative regulator of HIV-1 replication in CD4^+^ T-cells.

**Figure 2:**
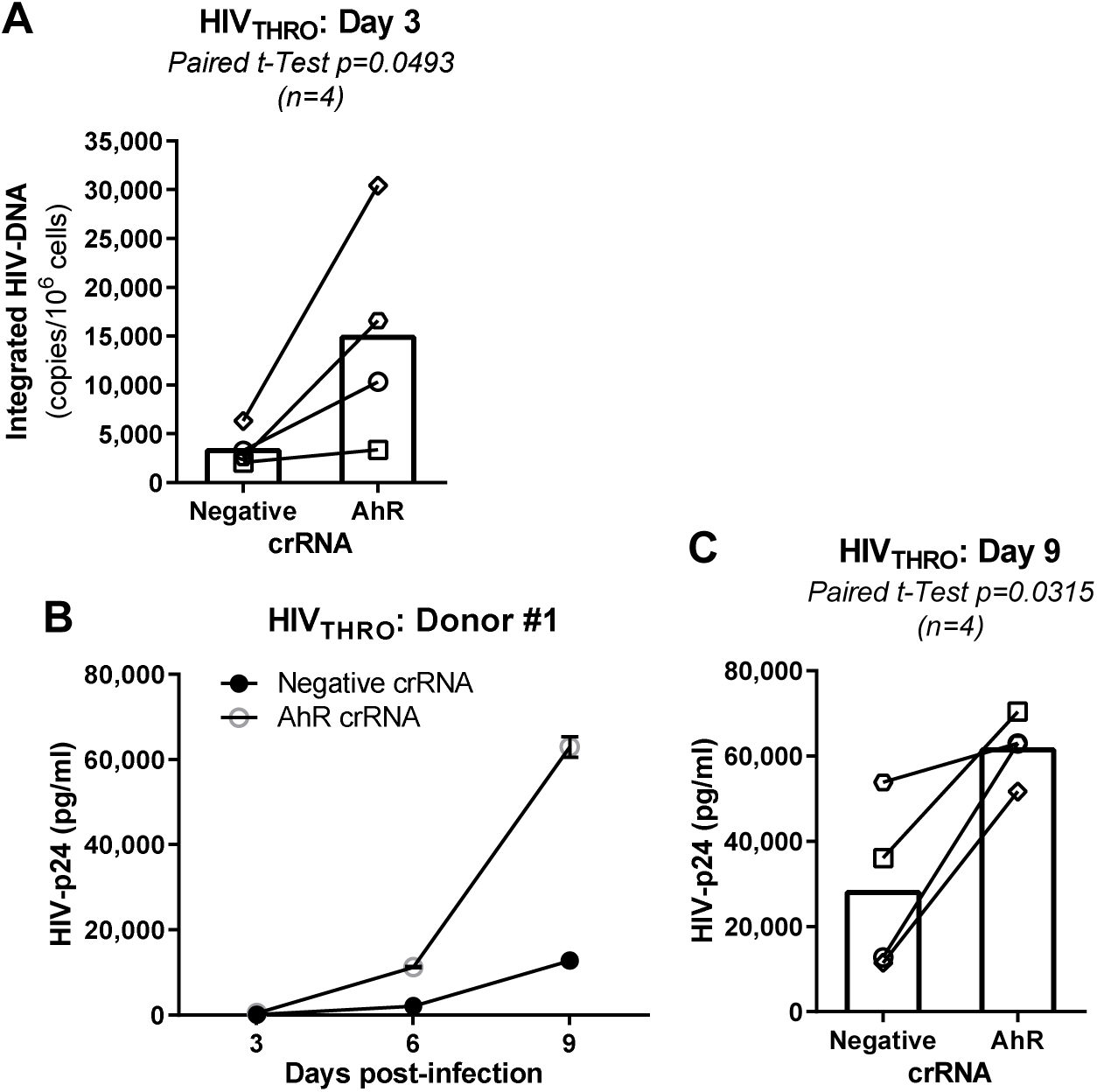
CRISPR/Cas9-mediated AhR KO renders primary CD4^+^ T-cells highly permissive to HIV-1 infection. Using the experimental flow chart depicted in Figure 1A, the effect of AhR KO on HIV-1 replication was investigated. Briefly, TCR-activated memory CD4^+^ T-cells were electroporated with AhR crRNA or negative crRNA in the presence of the tracrRNA. One day after electroporation, cells were exposed to HIV_THRO_ for 3 hours, the unbound virus was removed by extensive washing, and cells were cultured for 9 days in the presence of recombinant human IL-2 (5 ng/ml). Shown are levels of integrated HIV-DNA in cells harvested at day 3 post-infection (n=4) (**A**), as well as HIV-p24 levels in cell-culture supernatants at days 3, 6 and 9 post-infection in one representative donor **(B)** and statistical analysis of HIV-1 replication at day 9 post-infection in cells electroporated with Ahr crRNA *versus* negative crRNA (n=4) **(C)**. Paired t-Test p-values are indicated on the graph in panel **C**.

### Pharmacological AhR triggering/blockade modulates IL-22, IL-17A, IL-10 production and HIV-1 replication in memory CD4^+^ T-cells

AhR is a nuclear receptor with a transcriptional activity regulated by endogenous or exogenous ligands (32, 48). Thus, we further explored the effect of ligand-mediated AhR triggering *versus* blockade on Th-lineage cytokine production and HIV-1 replication. To do this, we used the AhR agonist 6-formylindolo [3,2-b]carbazole (FICZ), a photooxidation product of tryptophan (49), and the AhR antagonist CH-223191, a chemical inhibitor which blocks binding of ligands to AhR (50). As expected, the activation of memory CD4^+^ T-cells *via* the TCR in the presence of FICZ led to a >10-fold increase in CYP1A1 mRNA expression compared to DMSO, while exposure to CH223191 decreased CYP1A1 mRNA expression (<u>Supplemental Figure 4</u>). Subsequently, dose-response experiments demonstrated that CH-223191 decreased IL-22, IL-17A, and IL-10 but not IFN-γ production (<u>Supplemental Figure 5A-D, left panels</u>) at optimal doses (5 and 10 μM). At the opposite, FICZ increased IL-22 and IL-10, but not IL-17A and IFN-γ, production at 100 nM (<u>Supplemental Figure 5A-D, right panels</u>). At the concentrations tested, CH223191 and FICZ did not affect cell viability, nor cell proliferation estimated by the expression of Ki67 (<u>Supplemental Figure 6A-B</u>), a surrogate marker of the cell cyle S phase, thus excluding the cytotoxic effects of these drugs.

Further, to test the effects of AhR agonism/antagonism on HIV-1 replication, TCR-activated memory CD4^+^ T-cells cultured in the presence/absence of optimal CH223191 (10 µM) and FICZ (100 nM) doses were exposed to HIV_THRO_ and cultured in the presence/absence of CH223191 or FICZ for 9 days. CH223191 strongly promoted HIV-1 replication compared to DMSO, while FICZ induce a slight decrease (Figure 3A-B). Thus, pharmacological AhR blockade boosts HIV-1 replication in CD4^+^ T-cells, consistent with results generated in CRISPR/Cas9-mediated AhR KO (Figure 2).

**Figure 3:**
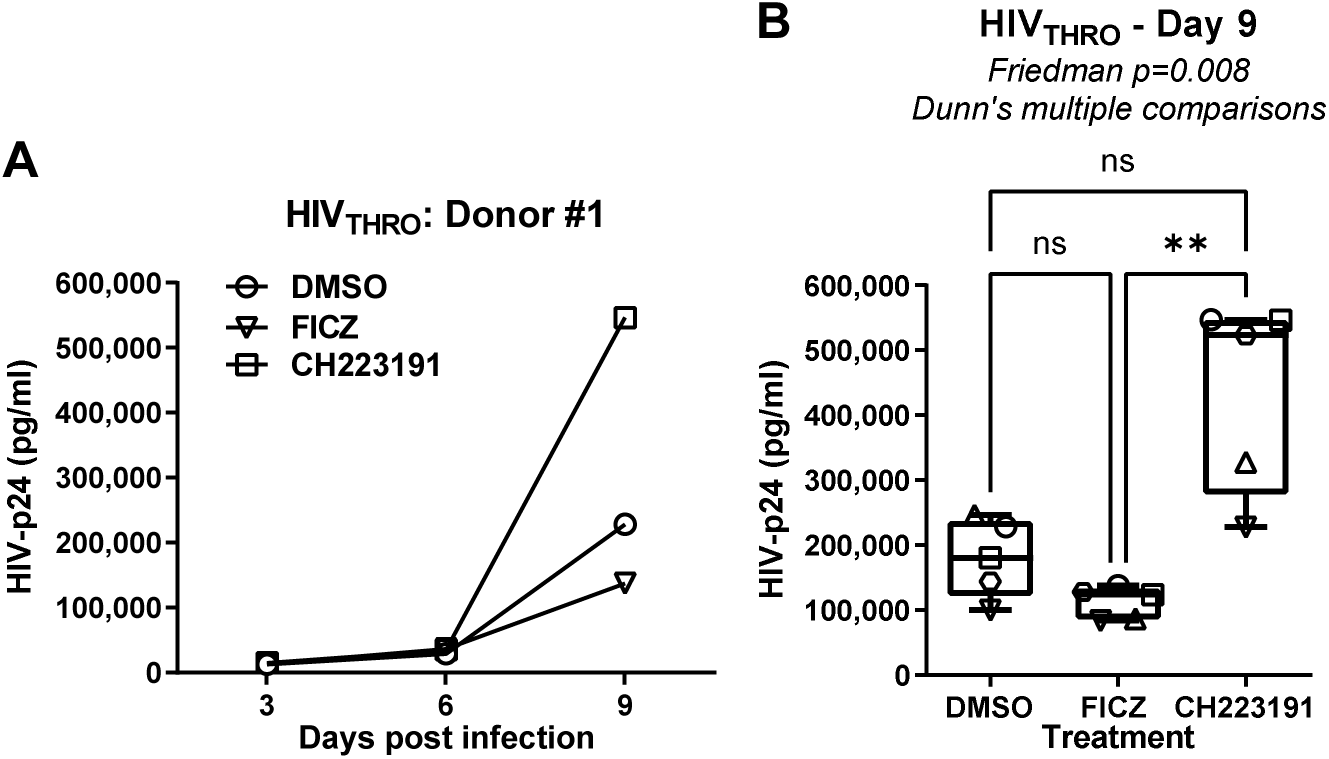
The AhR antagonism enhances T/F HIV THRO replication in memory CD4^+^ T-cells. Memory CD4^+^ T-cells isolated from PBMCs of HIV-uninfected individuals were activated *via* CD3/CD28 and cultured in the presence or the absence of the AhR antagonist CH223191 (10 µM) or agonist FICZ (100 nM) for 3 days, as in Figure 3. Cells were exposed to T/F HIV_THRO_ (50 ng HIV-p24/10^6^ cells) and cultured for up to 9 days in the presence of IL-2, in the presence or the absence of CH223191 or FICZ. Shown are HIV-p24 levels quantified by ELISA in cell-culture supernatants collected at days 3, 6, and 9 post-infection in one representative donor **(A)** and statistical analysis of results obtained using cells from n=5 donors, at day 9 post-infection **(B)**. Each symbol represents one donor. Shown are Friedman test p-values and Dunn’s multiple comparison significance (*, p < 0.05; **, p < 0.01; ***, p < 0.001).

It is noteworthy that biologically active doses of CH223191 and FICZ did not affect the expression of the HIV-1 receptor CD4, nor the co-receptors CCR5/CXCR4 (<u>Supplemental Figure 7A, left panels</u>). Similar to CRISPR-Cas9-mediated AhR KO (<u>Supplemental Figure 2</u>), CH223191 significantly downregulated ITGB7 expression, while FICZ tended to increase it (<u>Supplemental Figure 7D, left panels</u>).

Consistent with the documented antiviral role of IL-22 (51) and IL-10 (52, 53), recombinant IL-22 and/or IL-10 partially or completely counteracted the effect of CH223191 on HIV-1 replication in CD4^+^ T-cells (<u>Supplemental Figure 8</u>).

Together, these results provide evidence that AhR blockade boosts HIV-1 replication in CD4^+^ T-cells *via* mechanisms independent on CD4/CCR5/CXCR4 expression, but likely linked to the reduced production of the antiviral cytokines IL-22 and IL-10.

### Pharmacological AhR triggering/blockade modulates HIV-1 replication preferentially in CCR6^+^CD4^+^ T-cells

Among memory CD4^+^ T-cells, IL-17A and IL-22 are mainly produced by cells expressing the surface marker CCR6 (34, 40, 54). Also, CCR6^+^ compared to CCR6^-^ T-cells are reported to exhibit enhanced permissiveness to HIV-1 infection (8, 9, 13). Therefore, we sought to determine whether AhR modulates HIV-1 replication mainly in CCR6^+^ T-cells. To this aim, FACS-sorted memory CCR6^+^ and CCR6^-^ T-cells were stimulated *via* CD3/CD28 in the presence/absence of FICZ or CH223191 and exposed to HIV_THRO_. At day 3 post-TCR-triggering, prior infection, CCR6^+^ compared to CCR6^-^ T-cells preferentially produced IL-22 and IL-17A (Figure 4A-B). CH223191 compared to FICZ significantly decreased IL-22 and IL-17A production in CCR6^+^ T-cells (Figure 4C-D). Finally, CH223191 robustly increased HIV_THRO_ replication in CCR6^+^ T-cells, whereas the effects of AhR drugs on CCR6^-^ T-cells were only minor (Figure 4E-F). Collectively, these results demonstrate that the pharmacological AhR blockade facilitate HIV-1 replication in CCR6^+^ T-cells, while downregulating IL-22 and IL-17A production.

**Figure 4:**
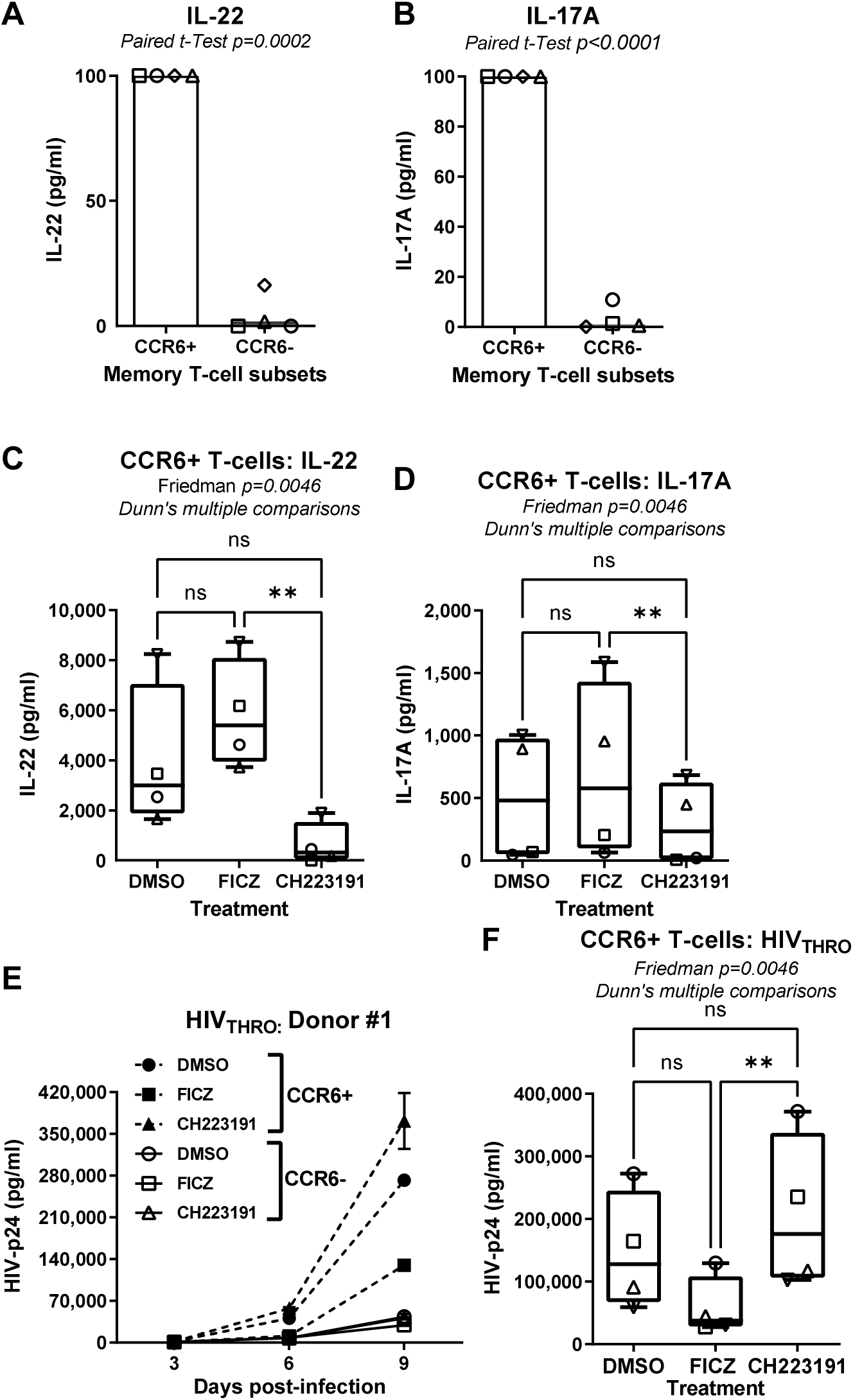
The AhR blockade increases HIV-1 replication in CCR6^+^ CD4^+^ T-cells. Flow cytometry-sorted memory CCR6^+^ and CCR6^-^ CD4^+^ T-cell subsets isolated from PBMCs of HIV-uninfected individuals were stimulated *via* CD3/CD28 Abs in the presence or the absence of the FICZ (100 nM) or CH223191 (10 µM) for 3 days. Cells were then exposed to T/F HIV_THRO_ (50 ng HIV-p24/10^6^ cells) and cultured in the presence of IL-2 and in the presence or the absence of AhR drugs. Levels of IL-22 **(A and C)** and IL-17A **(B and D)** were measured by ELISA in cell-culture supernatants of CCR6^+^ *versus* CCR6^-^T-cells (**A-B**) and CCR6^+^ T-cells treated or not with FICZ or CH223191 (**C-D**), at day 3 post-TCR triggering (n=4). Shown are HIV-p24 levels quantified by ELISA in cell-culture supernatant collected at day 3, 6 and 9 post-infection in one representative donor **(E)** and statistical analysis of results obtained using cells from n=4 different HIV-uninfected donors **(F)**. Each symbol represents 1 donor. Paired t-Test p-values (**A-B**) and Friedman test p-values and Dunn’s multiple comparison significance (**C-D and F**) are indicated on the graphs (*, p < 0.05; **, p < 0.01; ***, p < 0.001).

### Pharmacological AhR inhibition facilitates HIV-1 replication at early post-entry steps

To determine at which step of the viral replication cycle AhR ligands acted, TCR-activated memory CD4^+^ T-cells were exposed to single-round VSV-G-pseudotyped GFP-expressing HIV-1 (HIV_VSVG_-_GFP_), entering cells *via* the LDL receptor (55), independently of CD4 and co-receptors. CH223191 significantly increased cell-associated levels of early (RU5) and late (Gag) reverse transcripts and integrated HIV-DNA, when compared to DMSO or FICZ (Figure 5A-C). Consistently, increased levels of GFP expression, indicative of efficient HIV-1 transcription, were observed upon AhR blockade (Figure 5D-E). FICZ did not exert any significant effect in HIV-DNA levels or HIV-1 transcription when compared to DMSO (Figure 5A-E), likely due to the activation of the AhR pathway by ligands presents in the culture media (37). Indeed, TCR triggering leads to AhR pathway activation (<u>Supplemental Figure 1</u>). Together, our results demonstrate that pharmacological AhR blockade facilitates HIV-1 replication at early/late post-entry reverse transcription levels, subsequently leading to efficient viral integration, transcription and translation.

**Figure 5:**
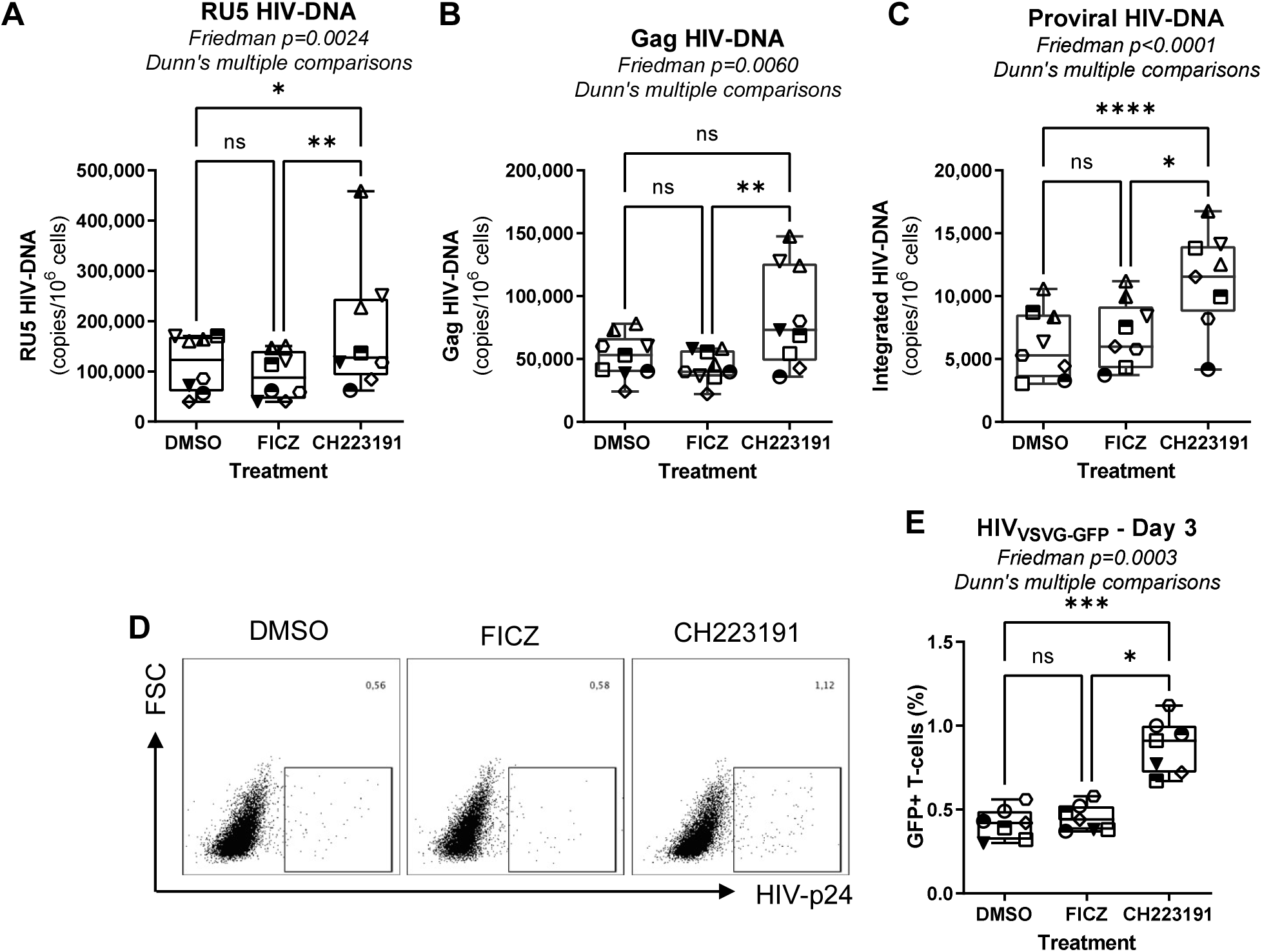
The AhR antagonism increases HIV-1 permissiveness in CD4^+^ T-cells at post-entry steps, between reverse transcription and integration. Memory CD4^+^ T-cells isolated from PBMCs of HIV-uninfected individuals were activated with CD3/CD28 Abs and cultured in the presence or the absence of CH223191 (10 µM) or FICZ (100 nM) for 3 days. Cells were then exposed to single-round VSV-G pseudotyped HIV-1 encoding *gfp* in place of *nef* (HIV_VSVG-GFP_) (50 ng HIV-p24/10^6^ cells) and cultured for 3 days in the presence or the absence of CH223191 or FICZ. Levels of early (RU5) **(A)** and late (Gag) reverse transcripts **(B)**, as well as integrated HIV-DNA levels **(C)**, were quantified by real-time nested PCR at day 3 post-infection. **(D-E)** The GFP expression was measured by flow cytometry as an indicator of HIV-1 transcription/translation. Shown is the frequency of GFP^+^ T-cells in one representative donor **(D)** and statistical analysis of GFP expression in T-cells from n=7 donors **(E)**. Each symbol represents one donor. **(A-C and E)** The Friedman test p-values and Dunn’s multiple comparison significance is indicated on the graphs (*, p < 0.05; **, p < 0.01; ***, p < 0.001).

### Pharmacological AhR inhibition boosts HIV-1 outgrowth in CD4^+^ T-cells of ART-treated PLWH

Quantitative viral outgrowth assays (QVOA) fail to predict HIV-1 rebound upon ART interruption, indicative of current limitations in the capacity to quantify replication-competent VR *in vitro* (6, 56). To explore the effect of AhR pathway on HIV-1 outgrowth, we used a simplified viral outgrowth assay (VOA) previously set up by our group (24, 57). Briefly, memory CD4^+^ T-cells from ART-treated PLWH (<u>Table 1, n=7</u>) were stimulated with CD3/CD28 Abs and cultured in the presence or the absence of AhR drugs, using the experimental procedure depicted in Figure 6A. In preliminary experiments, we demonstrated that TCR triggering in memory CD4^+^ T-cells of ART-treated PLWH results in AhR pathway activation, as demonstrated by the upregulation of CYP1A1 mRNA (<u>Supplemental Figure 9</u>). The AhR antagonist CH223191 significantly increased the frequency of HIV-p24^+^ T-cells (Figure 6B-C) and soluble HIV-p24 levels (Figure 6D), without affecting cell viability (Figure 6E). At the opposite, the AhR agonist FICZ showed only moderate effects on blocking HIV-1 outgrowth (Figure 6B-D), raising the possibility that this pathway may be already activated in cells from ART-treated PLWH.

**Figure 6:**
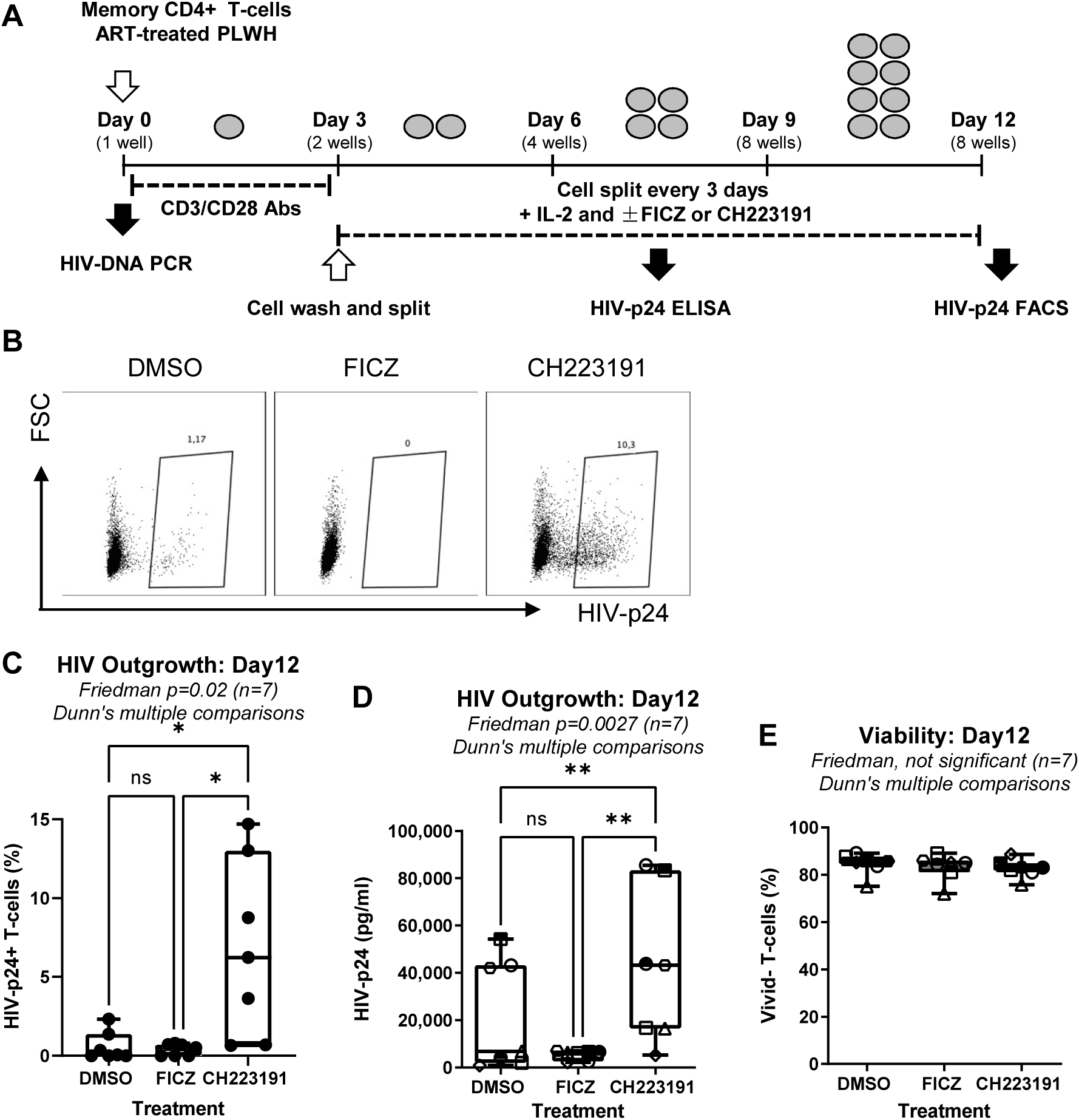
The AhR blockade boosts HIV-1 outgrowth in CD4^+^ T-cells of ART-treated PLWH. A viral outgrowth assay (VOA) was performed, as summarized in the experimental flowchart **(A)**. Briefly, memory CD4^+^ T-cells isolated from the PBMCs of ART-treated PLWH were activated with CD3/CD28 Abs and cultured in the presence of IL-2 and in the presence or the absence of FICZ (100 nM) or CH223191 (10 µM). At day 12 post-infection, HIV-1 replication and cell viability were measured. Show are intracellular HIV-p24 expression in T-cells (8 splitting replicates pooled per condition) from one representative donor (<u>ART #2;</u> Supplemental Table 1) **(B)**, as well as the statistical analysis of the % of HIV-p24^+^ cells **(C)**, HIV-p24 levels in cell-culture supernatants **(D)**, and the viability of cells harvested at day 12 of VOA **(E)** in experiments performed with cells form n=7 donors. Friedman test p-values, with Dunn’s multiple comparisons indicated on the graphs (*, p < 0.05; **, p < 0.01; ***, p < 0.001).

Taken together, these findings show that AhR blockade facilitates the detection of replication-competent VR in CD4^+^ T-cells of ART-treated PLWH.

### Genome-wide transcriptional profiling identifies new molecular mechanisms by which AhR blockade boosts HIV-1 outgrowth in CD4^+^ T-cells

To get insights into AhR-dependent regulatory mechanisms involved in viral outgrowth, RNA-Sequencing using the Illumina technology was performed on TCR-activated memory CD4^+^ T-cells of ART-treated PLWH cultured in the presence/absence of CH223191 (Figure 7A). Differentially expressed genes (DEG) were identified based on p-values, adjusted p-values, and fold change (FC) ratios (<u>Supplemental Files 1-2</u>). The volcano plot in Figure 7B illustrates DEG in response to CH223191 exposure. When DEG were analyzed based on adjusted p-values (<0.05) and ranked based on FC expression (cut-off >1.3), a number of 204 transcripts were upregulated (*e.g.,* CXCL13, CH25H, ALDH1A2, APOBEC3B, OASL, IL-27, CCR3, IL-6, CD38, CD276/B7-H3, IRF8, BACH2, DPP4, CD274/B7-H1/PD-L1, LAG3, IL-6R, NOD2, FASLG, TREML2, CXCL11, BATF3, PLAC8, AIM2, KLF7, CXCL10, HERC5, LGMN, IFI35, CD59, USP28, LEF1, TBX21, IRF1, MAPK13), while 258 were downregulated (*e.g.,* CYP1A1, CYP1B1, HIC1, AHRR, GPR15, GPR55, RARG, P2RY8, IL-22, ITGA1, CD226, ITGB7, CD109, TIPARP, CD160, IL-1a, TNFSF11A, TRAF4, IL-17F, UBD, LMNA, S1PR1, ITGAL, IL-1R1, RUNX2, IL-26, IL-24, CXCR6, ABCA2, ITGA4, TIGIT, PTGER2, CD27, NR4A2, RPTOR, CD96, LCK, CXCR3, FURIN, CXCR5, E2F7) (<u>Supplemental Files 1-2</u>). The downregulation of CYP1A1, CYP1B1, and AHRR (AhR repressor) by CH223191 validates the AhR-specificity of this antagonist (<u>Supplemental File 2</u>). Also, the downregulation of ITGB7 (<u>Supplemental File 2)</u>, was consistent with the results in <u>Supplemental Figures 2A-B and 7D</u>. Although changes in IL-10 mRNA did not reach statistical significance in RNA-Seq results, transcripts for IL-24, a cytokine reported to block the “*pathogenic*” functions of Th17 cells by inducing the expression of IL-10 (58), were found significantly downregulated by CH223191 (<u>Supplemental File 2</u>).

**Figure 7:**
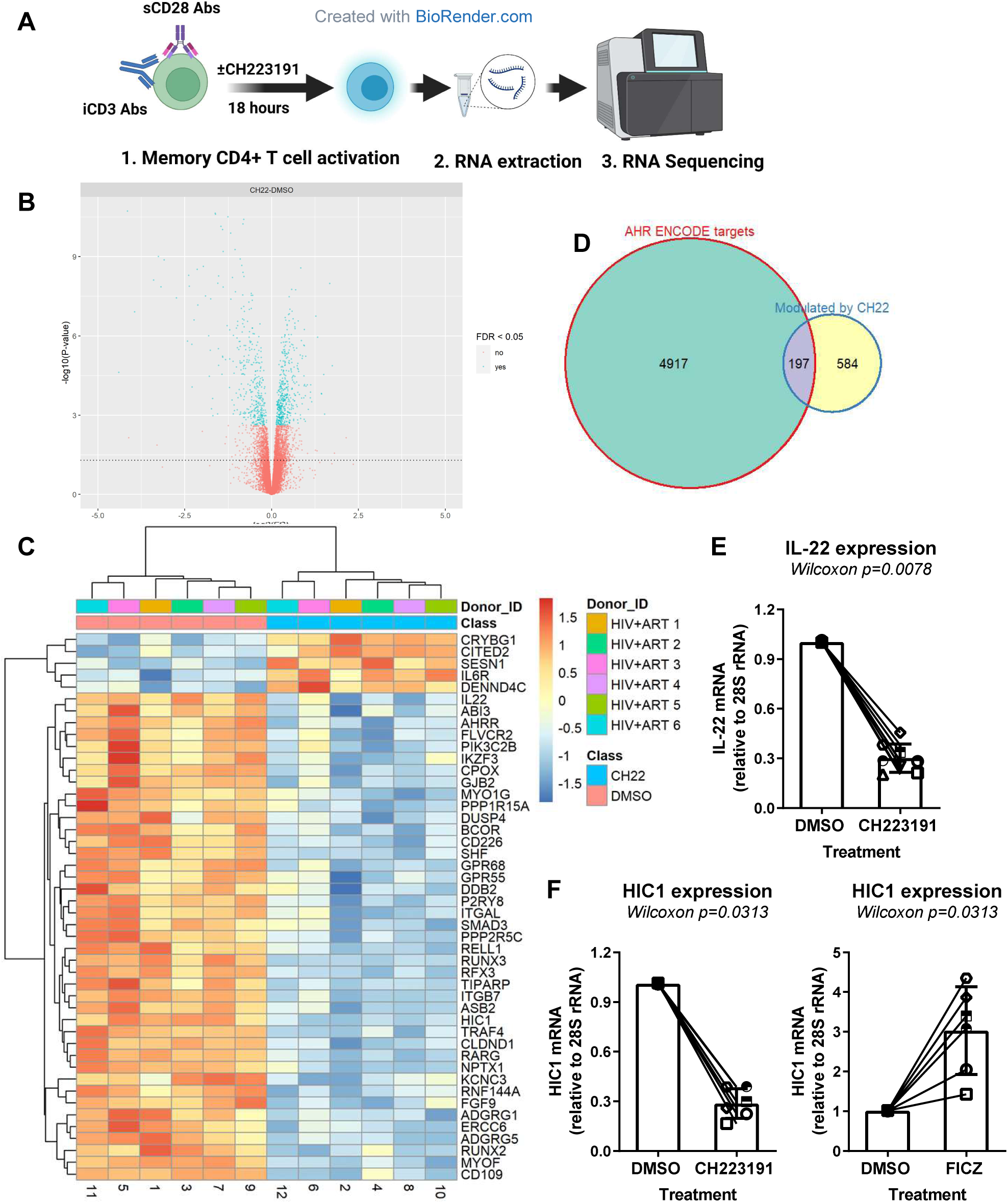
Transcriptional reprogramming of CD4^+^ T-cells upon pharmacological AhR inhibition. **(A)** Shown is the flow chart of RNA-Sequencing. Briefly, memory CD4^+^ T cells of ART-treated PLWH (n=6) were isolated and stimulated with CD3/CD28 Abs and cultured in the presence or the absence of CH223191 (10 µM) for 18 hours. Total RNA was extracted and used for RNA-Sequencing performed using the Illumina Technology. **(B)** Volcano plots for all probes in each linear model presented with the log2 FC on the x-axis and the negative logarithm of the adjusted P-values for false discovery rate (FDR) on the y-axis. The red/ blue color code is used based on the 5% FDR threshold. Shown are the representation of AhR direct target genes modulated by CH223191, as identified using the Encode Bioinformatic tool (https://www.encodeproject.org) **(C);**. top differentially expressed genes (DEG, identified based on p-values) in CD4^+^ T-cells exposed or not to CH22391 **(D)**.; IL-22 mRNA levels quantified by qPCR in CD4^+^ T-cells exposed or not to CH223191 (n=6) **(E)**; and HIC1 mRNA levels in CD4^+^ T-cells exposed or not to CH223191 or FICZ **(F)**. (**E-F)** Each symbol represents individual donor: bars represent median values. Wilcoxon matched-pairs signed rank test p-values are indicated on the graphs. **(G)** Finally, shown is our proposed model in which AhR regulates the expression of numerous genes, including HIC1, a transcriptional repressor documented to inhibit Tat-mediated HIV-1 transcription; it is noteworthy that CH223191 exposure results in decreased HIC1 expression, providing an explanation for the increased HIV-1 outgrowth upon AhR blockade.

Top 50 upregulated (*e.g.,* CRYBG1, CITED2, SESN1, IL-6R, DENND4C) and downregulated (*e.g.,* IL-22, AHRR, IKZF3, RELL1, RUNX3, ITGB7, RUNX2, HIC1) transcripts are depicted in Figure 7C. Among transcript modulated by CH223191, a fraction of them (197 out of 584 transcripts) were direct AhR transcriptional targets, as identified based on the expression of xenobiotic responsive elements (XRE) in their promoters using the Bioinformatics tool Encode (https://www.encodeproject.org) (Figure 7D). The list of genes modulated by CH223191, expressing XRE in their promoters include upregulated (*e.g.,* OASL, CCND1/cyclin D1, IL-6R, BATF3, SGK1, RORA, IFI35, LGALS3BP, CEBPB, GADD45A, and SIRT1) downregulated transcripts (*e.g.,* CYP1A1, SMAD3, TRAF4, IKZF3, IL1R1, RPTOR, and IL-32) (<u>Supplemental File 3</u>).

The downregulation of IL-22 mRNA levels upon CH223191 exposure (Figure 7C) was confirmed by RT-PCR (Figure 7E), thus validating the functional activity of AhR inhibition in these experimental settings

Among top downmodulated transcripts (Figure 7C; <u>Supplemental File 2</u>), the zinc-finger transcriptional repressor HIC1 (*Hypermethylated in cancer 1*) (59) was reported to inhibit Tat-dependent HIV-1 gene transcription (60) and to govern a unique transcriptional program involved in intestinal homeostasis (61) and T-cell tissue residency (62). Consistent with the RNA-Seq data, real-time RT-PCR quantification of HIC1 mRNA demonstrated that exposure to CH223191 significantly decreased, while exposure to FICZ robustly upregulated HIC1 mRNA expression (Figure 7F). Although the Encode search did not identify HIC1 as a direct AhR target (<u>Supplemental File 3</u>), *in silico* analysis using two other databases (*i.e,* Gene Cards, The human genome database and Eukaryotic promoter database) identified the presence of putative XRE in the HIC1 promoter (<u>data not shown</u>). Thus, HIC1 may be added to the list of new direct AhR gene targets (<u>Graphical abstract</u>).

To extract additional meaning from these RNA-Seq results, Gene set variation analysis (GSVA) was performed using the UC San Diego/Broad Institute Molecular Signature Database (MSigDB; https://www.gsea-msigdb.org/gsea/msigdb). This allowed the identification of top modulated canonical pathways in response to CH223191-exposed cells, including pathways upregulated (*e.g.,* T helper 1 differentiation, interferon alpha response, interferon gamma response, toll-like receptor 3 signaling, toll-like receptor 4 signaling, negative regulation of IL-2 production, toll-like receptor 9 signaling, positive regulation of IL-17 production, alpha beta T cell proliferation, and positive regulation of T cell proliferation) and downregulated (*e.g.,* zink ion homeostasis) (<u>Supplemental Figure 10A-B</u>). The modulation of these pathways may be either the direct effect of AhR-mediated transcriptional regulation and/or the consequence of HIV-1 sensing, consistent with the viral outgrowth promoted by CH223191 (Figure 6).

Further, data mining was performed by interrogating the NCBI HIV-1 interactor database (<u>Supplemental Figure 11A</u>). Among HIV-1 interactors, top upregulated transcripts included CD58, LGALS1, CD4, CCR3, DPP4, LGALS3BP, CEBPB, CD38, NFKBIA, PTK2B, HERC5, BACH2, SGK1, IL-6, NOTCH2, SNTB2, TNFRSF1A, CD59, SIRT1, CD28, and LDLR, while top downregulated transcripts included CD27, LCK, ITGB7, CD226, ITGAL, IKZF3, IL-1, GPR15, ITGA4, CXCR6, and CXCR3 (<u>Supplemental Figure 11A</u>). Real-time validations were performed for SIRT1, CEBPB, and BACH2 mRNA. Results demonstrated that CH223191 only slightly modified SIRT1 mRNA expression, while FICZ significantly decreased it (<u>Supplemental Figure 12A</u>). There was a tendency for increased and decreased CEBPB mRNA expression upon exposure to CH223191 (p=0.219) and FICZ (p=0.07), respectively, but differences did not reach statistical significance (<u>Supplemental Figure 12B</u>). Finally, CH223191 increased (p=0.0525), while FICZ decreased (p=0.0062) BACH2 mRNA expression in TCR-activated memory CD4^+^ T-cells (<u>Supplemental Figure 12C</u>). Of note, SIRT1, CEBPB, and BACH2 are included on the list of putative AhR targets (<u>Supplemental File 3</u>).

Together, these results revealed the transcriptional reprogramming associated with CH223191-mediated increase in HIV-1 outgrowth in CD4^+^ T-cells of ART-treated PLWH and identified HIC1, SIRT1, CEBPB and BACH2 as novel AhR transcriptional targets and HIV interactors (Figure 8).

## Discussion

In this manuscript, we provide evidence that AhR activation subsequent to TCR triggering acts as a barrier to HIV-1 infection and outgrowth in CD4^+^ T-cells. The genetic and pharmacological inhibition of AhR promoted increased HIV-1 replication *in vitro*. Pharmacological inhibition of AhR also enhanced viral outgrowth from CD4^+^ T-cells of ART-treated PLWH. RNA-Sequencing revealed the transcriptional reprogramming associated with increased viral outgrowth prompted by AhR blockade, with *in sillico* analyses identifying novel putative AhR target genes listed on the NCBI HIV-1 Interactor Database (Figure 8). Together, these results support a model in which AhR activation plays a dual role during HIV-1 infection: first, AhR activation may limit HIV-1 acquisition during primary infection in CD4^+^ T cells and second, AhR governs a transcriptional program that potentially maintain “*silent*” VR in ART-treated PLWH. Indeed, a state of deep latency was observed in CD4^+^ T-cells residing in the gut (63–65), an anatomic site rich in AhR ligands (30–32). In the context where AhR antagonists are currently tested in clinical trials for cancer (44, 45), their ability to reactivated and purge VR in ART-treated PLWH deserves investigations in “*shock and kill*” strategies.

TCR triggering in CD4^+^ T-cells is associated with an increased permissiveness to HIV-1 replication (66), in part, as a consequence of a reduced expression/function of HIV-1 restriction factors, such as SAMHD1 (67). One important finding of our study is the demonstration that AhR expression at RNA/protein levels, along with its activity (CYP1A1 transcription), are upregulated upon TCR triggering. This supports the idea that the AhR expression/activity prevents excessive HIV-1 replication in cognate activated CD4^+^ T-cells. We demonstrated that AhR pathway controls HIV-1 replication at post-entry levels, as indicated by no changes in the expression of the HIV-1 receptor CD4 and co-receptors CCR5/CXCR4. Also, single-round infection with VSV-G-pseudotyped HIV-1 (HIV_VSVG-GFP_) revealed that CH223191-mediated AhR blockade increased the efficacy of early/late HIV-1 reverse transcription, subsequently favoring HIV-DNA integration and *de novo* viral protein expression. Finally, CH223191 strongly boosted HIV-1 outgrowth from CD4^+^ T-cells of ART-treated PLWH. Our results are consistent with previous studies by Kueck *et al.* reporting an antiviral role for AhR in macrophages (43). In the latter study, by using the FICZ and CH223191, the authors demonstrated that AhR activation restricts productive HIV-1 replication *via* the regulation of cyclin-dependent kinase (CDK)1/2 expression and SAMHD1-mediated restriction of HIV-1 reverse transcription (43). Although our RNA-Sequencing results did not reveal the modulation of CDK1/2 nor SAMHD1 mRNA expression in CD4^+^ T-cells, our RNA-Sequencing data revealed an upregulation of cyclin D1 (CCND1) by CH223191. The involvement of AhR in controlling the dNTP levels *via* SAMHD1 phosphorylation/dephosphorylation in CD4^+^ T-cells remains a possibility to be tested.

The proviral effects of CH223191 were mainly observed in Th17/Th22-polarized CCR6^+^ T-cells, consistent with the fact that Th17 cells preferentially express AhR (34, 37) and exhibit increased permissiveness to HIV-1 infection (13). AhR regulates the expression of multiple downstream effector genes like CYP1A1 and CYP1B1 (32, 33). Also, AhR was reported to transcriptionally regulate expression of IL-22, IL-17A, and IL-10 in CD4^+^ T-cells (25, 27, 34, 37, 41). While IL-17A exhibit proviral features (68), the antiviral effects of IL-22 and IL-10 are documented (51, 52), with IL-10 contributing to VR persistence in SIV infection models (53). Therefore, the downregulation of IL-22 and IL-10 by CH223191 may explain, at least in part, the increased viral outgrowth observed in CD4^+^ T-cells of ART-treated PLWH. Consistent with the fact that AhR is a marker for “*non-pathogenic*” Th17 cells, the pharmacological inhibition of AhR increased the expression of the “*pathogenic*” Th17 markers IL-23R (25), SGK1 (69) and RORA (70).

Of particular importance, our results revealed that, similar to macrophages (42), the expression of the gut-homing molecule ITGB7 is also under the control of AhR in CD4^+^ T-cells. Studies by our group demonstrated that ITGB7 is a marker for CCR6^+^CD4^+^ T-cells highly permissive to HIV-1 infection (71). In addition to the fact that ITGB7 binds HIV-gp120 (72), the interaction of ITGB7 with its ligand MadCAM-1 further boosts HIV-1 permissiveness in ITGB7^+^ T-cells (73). Thus, the transcriptional program governed by AhR may imprint CD4^+^ T-cells with a gut-homing tropism, thus facilitating their HIV-1 infection at portal sites of viral entry. In addition to ITGB7, other transcripts for chemokines, chemokine receptors, and adhesion molecules involved in tissue-specific trafficking were upregulated (CCR3, CXCL13, CXCL11, CXCL10) or downregulated (GPR15, ITGA1, CD226, ITGA2B, S1PR1, ITGAL, ITGA4, CXCR6, CXCR3, CXCR5) by CH223191. GPR15 mediate gut-homing under the control of AhR (74), S1PR1 is a master regulator of T-cell egress from the lymph nodes (75), while CCR3 was previously documented by our group to be preferential expressed on HIV-infected cells *in vitro* (76). Of particular notice, HIC1, strongly induced by FICZ and decreased by CH223191 in CD4^+^ T-cells of ART-treated PLWH, represents a key regulator in intestinal homeostasis (61) and was recently reported to promote T-cell tissue residency (62). Future studies should explore the role of the AhR-HIC1 interplay in controlling the tissue-residency in the gut, mainly during HIV-1 infection, a condition associated with increased expression of AhR ligands (77, 78). Based on our results, one may anticipate that AhR blockade promotes the recirculation of tissue-residence CD4+ T-cells carrying VR (Figure 8).

The RNA-Sequencing performed on bulk memory CD4^+^ T-cells revealed that CH223191 downregulated the expression of CYP1A1, AHRR, and TIPARP, involved in the negative feedback regulation of AhR signaling (32, 33), indicative of a diminished AhR transcriptional activity. In addition, we observed the modulation of other lineage cytokines important for HIV-1 pathogenesis, including the downregulation of IL-1α, IL-1R1, IL-24 and IL-26 and the upregulation of IL-27, IL-6, IL-6R, IL-7R, IL-18RAP, and IL-12RB1 expression. Among these transcripts, the ENCODE search revealed that IL-6R and IL-1R1 are putatively directly regulated by AhR, as they express XRE in their promoters. Thus, AhR activation may facilitate T-cell responsiveness to IL-1 to the detriment of IL-6. Although IL-27 was reported to inhibit HIV-1 replication in macrophages (79), its expression appears compatible with efficient HIV-1 replication in CD4^+^ T-cells exposed to CH223191. Furthermore, we report here that genes encoding for immune check point activators/inhibitors modulated by AhR, as reflected by the upregulation of CD276/B-H3, CD38, CD274/PD-L1, LAG3, and CD59, and the downregulation of CD109, CD160, TIGIT, CD96, and CD27 in the presence of CH223191. Whether these AhR-modulated transcripts represent novel surface markers for T-cells carrying replication-competent VR remains to be evaluated using single-cell analysis techniques. However, multiple DEG did not express putative AhR binding elements in their promoter; thus, their modulation may be either the results of an indirect regulation or the consequence of HIV-1 sensing, as it is the case of the IFN-induced genes CXCL10, IRF8, IFI35, and IRF1 that are modulated by CH223191. Nevertheless, the ENCODE database may not be complete and chromatin immunoprecipitation studies are required to properly identify AhR target genes in human CD4^+^ T-cells.

In TCR-activated CD4^+^ T-cells of ART-treated PLWH, CH223191 modulated the expression of various sets of genes previously identified as HIV interactors. Among them, HIC1 was documented to inhibit Tat-dependent HIV-1 transcription (60), raising the possibility that HIC1 downregulation is one mechanisms of by which viral outgrowth was boosted by CH223191. This is consistent with our *in sillico* finding that AhR binding elements are present in the HIC1 promoter, suggesting a direct regulation of HIC1 transcription by AhR (<u>Graphical abstract</u>). Of note, the down regulation of HIC1 coincided with the upregulation of IL-6, a pro-inflammatory cytokine associated with HIV-1 disease progression (80) and HIV-1 replication (52) and a negative regulator of HIC1 expression (81). In addition, SIRT1, CEBPB, and BACH2 were also identified as HIV interactors included on the list of putative AhR gene targets. Of note, SIRT1 (sirtuin 1) is a histone deacetylase repressed at transcriptional level by HIC1 and which, in turn, regulates HIC1 activity (59, 60). Also, SIRT1 facilitates Tat-dependent HIV-1 transcription (82) and represents a therapeutic target also in HIV-1 infection (83). SIRT1 is used as a therapeutic target in cancer (84). CEBPB (CCAAT/enhancer-binding protein beta), involved in the transcriptional regulation of multiple genes, including HIV-1 (85). Finally, BACH2 caught our attention based on reports demonstrating preferential HIV-1 integration in this gene (86–88). These results raise the possibility that CH223191-mediated BACH2 expression may facilitate HIV-1 integration into this gene; alternatively, since HIV-1 preferentially integrates in BACH2 (86–88), CH223191-mediated BACH2 transcription may explain the increased viral outgrowth.

The evidence in this manuscript that AhR acts as a barrier to HIV-1 infection/outgrowth in TCR-activated CD4^+^ T-cells, is in line with studies by Kueck et al. performed in macrophages (43). In contrast, a report by Zhou *et al.* demonstrated that exposure to AhR ligands (*e.g.,* kynurenine, FICZ) promoted viral transcription in resting CD4^+^ T-cells of ART-treated PLWH and reported the presence of AhR-binding XRE in the HIV-1 promoter (89). These controversial results raise new questions on potentially different AhR-mediated mechanisms of action in resting *versus* TCR-activated CD4^+^ T-cells. Also, the capacity of AhR to bind directly to the HIV-1 promoter as an activator *versus* a repressor may dependent on the cell activation status and the expression of other HIV-1 dependency factors. These aspects remain to be clarified.

In addition to its role as a ligand-activated transcription factor, AhR exhibit an intrinsic E3 ubiquitin ligase activity that targets the degradation of steroid receptors (*e.g.,* estrogen receptors, ER) (90). Although this alternative function of AhR was not explored in this study, the interaction between AhR and ER deserves further investigation in the context where ER were reported to inhibit HIV-1 transcription (91). By degrading ER, AhR may facilitate HIV-1 reactivation from latency. The AhR switch from a transcription factor (genomic action) to an E3 ubiquitin ligase (non-genomic action) depends on ARNT and AHRR (92). Molecular mechanisms by which genomic *versus* non-genomic AhR-dependent regulatory pathways are triggered remain to be elucidated, and may depend on the target cells, as well as on the type of ligand present in the cellular environment, its concentration and the duration of exposure (32). In our study, we did not differentiate between genomic and non-genomic actions of AhR-mediated functions in CD4^+^ T-cells; also, we did not perform experiments in CD4^+^ T-cells from males and females to determine the influence of sex on AhR-mediated regulation of HIV-1 replication in CD4^+^ T-cells. Future studies should address these issues, in an effort to explain controversies on the role of AhR in antiviral immunity.

Altogether, results included in this manuscript support a model in which AhR transcriptionally reprograms Th17/Th22-polarized CD4^+^ T-cells toward the expression of tissue residence markers (*e.g.,* ITGB7, CXCR6) and acts as a barrier to HIV-1 infection and viral outgrowth (Figure 8). In this model, one may anticipate that in uninfected individuals (in the presence of natural non-toxic AhR ligands) AhR activation in Th17/Th22 cells may strengthen the integrity of the intestinal barrier and create a state of viral refractoriness that prevents mucosal HIV-1 transmission. At the opposite, in PLWH (in the presence of toxic AhR ligands) AhR activation may facilitate the persistence of latent HIV-1 reservoirs during ART. This model is consistent with previous studies demonstrating the abundance of AhR ligands in the GALT (30–32), the abnormal expression of IDO-1 and Trp catabolites in ART-treated PLWH (77, 78), and the maintenance of a state of deep HIV-1 latency at intestinal level (63–65). Whether VR latency in ART-treated PLWH correlate with the predominance of deleterious AhR ligands remains to be determined. Future studies are needed to identify AhR antagonists, such as those currently tested in clinical trials for cancer (44, 45), that may be used in an effort to purge HIV-1 reservoirs in ART-treated PLWH.

## MATERIALS AND METHODS

### Subjects

Leukapheresis were collected from ART-treated PLWH and HIV-uninfected study participants recruited at the McGill University Health Centre and Centre Hospitalier de l’Université de Montréal (CHUM, Montreal, Québec, Canada). Peripheral blood mononuclear cells (PBMCs) (10^9^–10^10^) were isolated from leukapheresis and cryopreserved as previously described (93). The clinical characteristics of ART-treated PLWH are listed in Supplemental Table 1.

### Ethics statement

In this study, leukapheresis collection from ART-treated PLWH and HIV-uninfected study participants was performed in compliance with the principles included in the Declaration of Helsinki. This study is approved by Institutional Review Board of the McGill University Health Centre and the CHUM-Research Centre, Montreal, Quebec, Canada. All study participants have provided signed informed consents and agreed to publish the results in Journal with their samples.

### Flow-cytometry analysis

For flow cytometry analysis antibodies (Abs) were listed in <u>Supplemental Table2</u>. Intracellular staining was performed with BD Cytofix/Cytoperm kit (BD Biosciences, Franklin Lakes, NJ, USA), as we previously described (68, 94). The viability dye LIVE/DEAD Fixable Aqua Dead Cell Stain Kit (Vivid, Life Technologies, CA, USA) was used to exclude dead cells. Flow cytometry analysis was performed using a LSRII cytometer (BD Pharmingen San Diego, CA, USA). The positivity gates were placed using the fluorescence minus one (FMO) strategy. All flow cytometry data were analyzed with the FlowJo software version 10.6.1 (Tree Star, Inc).

### Cell sorting and TCR activation

Memory CD4^+^ T cells with (>95% purity) were enriched from PBMCs by negative selection using magnetic beads (Miltenyi, Bergisch Gladbach, Germany) (68, 93, 94). In other experiments, CCR6^+^ and CCR6^-^ memory CD4^+^ T-cells were sorted by flow cytometry (BDAria III), as we previously described (93, 94). Cells were activated using immobilized CD3 and soluble CD28 antibodies (1 µg/mL; BD Pharmingen San Diego, CA, USA) for 3 days and used for subsequent experiments.

### AhR ligands

The following AhR drugs were used in this study: the antagonist CH-223191 (Sigma, St. Louis, MO, USA; concentrations: 1.25, 2.5, 5, and 10 µM) and the agonist FICZ (Sigma, St. Louis, MO, USA; concentrations: 10 and 100 nM). Stock solutions of these ligands were prepared in dimethylsulfoxide (DMSO), aliquoted and stored according to the manufacturer’s instructions.

### HIV-1 infection *in vitro*

The following HIV-1 strains were used: replication-competent CCR5-tropic Transmitted Founder T/F THRO (HIV_THRO_) and NL4.3BaL (HIV_NL4.3BaL_), and single-round VSVG-HIV-GFP (HIV_VSVG_). The HIV_THRO_ molecular clone plasmid (Catalog No. 11919) was obtained from Dr. John Kappes and Dr. Christina Ochsenbauer through the NIH AIDS Reagent Program, Division of AIDS, NIAID, NIH (95). HIV_NL4.3BaL_ is a NL4.3-based provirus expressing the BaL envelope, provided by Michel Tremblay (U Laval, Québec) and originating from Dr Roger J Pomerantz (Thomas Jefferson University, Philadelphia, Pennsylvania, USA). The HIV_VSVG_ stock was generated using a plasmid encoding for an env-NL4.3-based provirus, with *gfp* in place of *nef*, and another plasmid encoding for VSV-G, as previously described (93, 96). HIV-1 stocks were generated by Fugene-mediated transection, as we previously described (93, 96). Memory CD4^+^ T-cells activated *via* CD3/CD28 for 3 days were exposed to HIV-1 (25 ng HIV-p24/10^6^ cells) for 3 hours at 37 °C. Cells were washed to remove the unbound virions. Then, cells were cultured in the presence of IL-2 (5 ng/ml; R&D Systems, Minneapolis, MN, USA), and in the presence or the absence of CH-223191 or FICZ, at the indicated concentrations up to 9 days, with media being refreshed every 3 days. Viral replication was monitored by flow cytometry upon HIV-p24 intracellular staining and by ELISA soluble HIV-p24 quantification, as previously described (68, 93, 96). In parallel, HIV-infected cells were harvested at day 3 post-infection for PCR-based HIV-DNA quantification as described below.

### Quantification of Gag and integrated HIV-DNA

RU5 and Gag reverse transcripts and integrated HIV-DNA levels were quantified by nested real-time PCR (Light Cycler 480, Roche). Experiments were performed in triplicates on 0.5-1×10^5^ T-cells/PCR. HIV-1 copy numbers were normalized to CD3 copy numbers (two CD3 copies per cell), as previously described (93, 96).

### Viral outgrowth assay (VOA)

The VOA was performed, as we previously described (57, 94). Briefly, memory CD4^+^ T-cells (1×10^6^ cells/ml RPMI, 10% FBS, 1% Penicillin/Streptomycin) were isolated by MACS from PBMCs of ART-treated PLWH and cultured in 48-well plates in the presence of CD3/CD28 Abs (1 µg/ml). At day 3 post-TCR triggering, cells were washed and split into two new 48-well plate wells and cultured with IL-2 (5 ng/ml; R&D Systems), in the presence or the absence of CH-223191 (10 µM) and FICZ (100 nM). Cells were further split at day 6 and 9 post-stimulation and half of the media was replenished with IL-2 containing or not AhR drugs. Cells were harvested at day 12 and the intracellular HIV-p24 expression was measured by flow-cytometry. In parallel, soluble HIV-p24 levels were quantified by ELISA in the collected supernatant of days 3, 6, 9 and 12, as previously described (93, 96).

### CRISPR/Cas9-mediated AhR KO

To KO AhR expression, we used the CRISPR/Cas9 technology, and a published protocol adapted for primary CD4^+^ T-cells (46) (Figure 1A). Briefly, memory CD4^+^ T-cells were stimulated *via* CD3/CD28 for 3 days prior to electroporation. Electroporation was performed using the Amaxa P3 Primary Cell 96-well Nucleofector kit and 4D-Nucleofecter (Lonza, Walkersville, MD, USA). crRNAs were selected from CRISPR sgRNA database of (Genscript, USA). crRNA, tracrRNA and Cas9 were purchased from IDT (IDT, Newark, NJ, USA). crRNA and *trans*-activating crRNA (tracrRNA) were mixed at equimolar concentrations in a sterile PCR tube. Oligos were annealed by heating at 95°C for 5 min in PCR thermocycler and the mix was slowly cooled to room temperature. An equimolar concentration of Cas9-NLS was slowly added to the crRNA:tracrRNA and incubated for 10 minutes at room temperature to generate Cas9 RNPs complex. For each reaction, 1 x 10^6^ T-cells were pelleted and re-suspended in 100 μL nucleofection solution (Lonza, Walkersville, MD, USA). T-cell suspensions were mixed with 5 μl RNP complex and cell/RNP mix transferred to Nucleofector cuvette. Cells were electroporated using program T-020 on the Amaxa 4D-Nucleofector (Lonza, Walkersville, MD, USA). After nucleofection, pre-warmed T-cell media was used to transfer transfected cells in 48-well plates and cells were allowed to culture in the presence of IL-2 (5 ng/ml) for 3-5 days. The AhR gene KO was confirmed by Western Blot (Figure 1B). crRNA sequences were as follows: AhR#1: 5’-AAGTCGGTCTCTATGCCGCT −3’ and AhR#2: 5’-TTGCTGCTCTACAGTTATCC −3’.

### T7 endonuclease assay

The AhR gene KO was also confirmed by T7 endonuclease assay, as previously described (97). Cells were lysed with DNAzol (Invitrogen, Waltham, Massachusetts, USA) and genomic DNA was extracted. AhR target regions were PCR-amplified using the forward primer 5’-GGGAATCACTGTGCTACAAATGC-3’and the reverse primer 5’-CAGAAAATCCAGCAAGATGGTGT −3’. The amplification was performed using the PFU HiFi DNA polymerase (NEB, Germany) following the manufacturer’s instructions. The PCR products were gel-purified using the QIAEX II Gel extraction kit (Qiagen, Germany). After purification, the PCR products were denatured and slowly re annealed to form hetero-duplex DNA and digested with T7 endonuclease (NEB, Germany) for 30 min at 37 °C. The resulting DNA fragments were visualized by 1.5% agarose gel electrophoresis.

### AhR siRNA

AhR RNA interference were performed in primary CD4^+^ T cells, as described previously (98). Briefly, memory CD4^+^ T-cells were stimulated by CD3/CD28 Abs for 2 days. Activated cells were nuclofected with 100 µM AhR or non-targeting (NT1) siRNA (ON-TARGETplus SMART pool, Dharmacon) using the Amaxa Human T cell Nucleofector Kit (Lonza, Walkersville, MD, USA), according to the manufacturer’s protocol. Nucleofected cells (2×10^6^) were cultured for 24 h at 37 °C in the presence of IL-2 (5 ng/ml). Cells were exposed to HIV-1 and cultured up to 3 days. To check the efficacy of KO, the AhR gene expression level was assessed by SYBR Green real-time RT-PCR 24h post-nucleofection. At day 3 post-infection, cells were stained with LIVE/DEAD® Fixable Dead Cell Stain Kit (Vivid, Life Technologies,CA) were analyzed by FACS (BD LSRII).

### SYBR Green quantitative real time RT-PCR

Total RNA was isolated using the RNAeasy mini kit (Qiagen, Hilden, Germany). The AhR mRNA was quantified by one step SYBR Green real-time RT-PCR using the AhR Quantitect primer (Qiagen, Hilden, Germany). The RT-PCR reactions were carried out using the Light Cycler 480 II (Roche, Basel, Switzerland). The relative expression of AhR was normalized to the house keeping gene 28S rRNA. Each reaction was performed in triplicates.

### Western Blot

Cells were lysed with the RIPA buffer (Cell Signaling) supplemented with phosphatase inhibitors (PhosSTOP) and protease inhibitor cocktail (Complete Mini EDTA-free, Roche) for 10 minutes at 4 °C and centrifuged at 14,000 g for 10 minutes. Proteins were quantified by Bradford assay (Bio-Rad, Hercules, CA, USA). Samples (15-30 µg protein/well) were loaded on SDS PAGE gel (90 minutes, 100 V), transferred to PVDF membrane (Millipore) and blocked with 5% BSA. Subsequently membranes were incubated with primary AhR antibodies (Clone ab190797, Abcam, Cambridge, UK) and β-actin (Sigma, USA). Immunoreactive bands were detected using HRP-conjugated secondary antibodies and revealed with ECL substrate (GE Healthcare, Chicago, IL, USA).

### ELISA

HIV-p24 levels were quantified in cell culture supernatants using a homemade ELISA, as described previously (93). IL-17A, IL-10 and IL-22 (DuoSet ELISA R&D Systems, Minneapolis, MN, USA) IFN-γ (Invitrogen, Waltham, Massachusetts, USA) cytokine levels in cell culture supernatants were measured, as per manufacturer’s instructions.

### Illumina RNA Sequencing and analysis

CD4^+^ memory T-cells were isolated from ART-treated PLWH, activated with CD3/CD28 Abs, and cultured in the presence or the absence of the AHR antagonist CH223191 (10 µM) for 18h.Total RNA was extracted using RNeasy Plus mini kit (Qiagen, Germantown, Maryland, USA). Genome-wide transcriptional profiling was performed by Genome Québec (Montreal, Québec, Canada) using the Illumina RNA-Sequencing technology (NovaSeq6000 S4 PE 100bp 25M reads). Briefly, the paired-end sequencing reads were aligned to coding and non-coding transcripts from Homo Sapiens database GRCh 37 version75 and quantified with the Kallisto software version 0.44.0. The entire RNA-Sequencing data set and the technical information requested by Minimum Information About a Microarray Experiment (MIAME) are available at the Gene Expression Omnibus database under accession **<u>GSE198078</u>**.

Differentially expressed genes (DEG) were identified based on p-values (p<0.05), adjusted p-values (adj. p<0.05) and fold-change (FC, cutoff 1.3). Statistical analyses were performed using R version 4.21. Differential expression analysis was performed using the limma Bioconductor R package (version 3.52.2) on the log2-counts per million (logCPM) transformed transcript-level and gene-level data. Gene set enrichment analysis (GSEA; C2, C3, C5, C7, C8, and Hallmark databases) was performed using the GSVA method (package version 1.344.2) on the logCPM data using a Gaussian cumulative distribution function. Finally, ChIP-seq peaks annotation of AhR targets (ENCODE dataset ENCFF763XLN; reference genome HG19) was done using packages ChIPseeker (version 1.32.1) and ENCODExplorerData (version 0.99.5).

### Statistical analysis

Statistical analyses were performed using the Prism 7 (GraphPad, Inc., La Joya, CA, USA) software. In each graph and Figure legend specific statistical test were applied. P-values are indicated on the graphs with statistical significance as follows*P < 0.05; **P < 0.01; ***P < 0.001; ****P < 0.0001

## Acknowledgement

The authors thank Dr. Dominique Gauchat and Philippe St Onge (Flow Cytometry Core Facility, CHUM-Research Center, Montréal, QC, Canada) for expert technical support with polychromatic flow cytometry sorting; Olfa Debbeche (Biosafety Level 3 Core Facility CHUM-Research Center, Montréal, QC, Canada); Mario Legault (FRQ-S/AIDS and Infectious Diseases Network; Montré al, QC, Canada) for help with ethical approvals and informed consents; Josée Girouard and Angi e Massicotte (McGill University Health Centre, Montréal, QC, Canada) for their key contribution to blood collection and clinical information from PLWH and uninfected study participants. The authors also thank Dr Roger J Pomerantz (Thomas Jefferson University, Philadelphia, Pennsylvania, USA) and Dr Michel Tremblay (Université Laval, Quebec, QC, Canada) for providing us with the NL4.3BaL pseudotyped HIV. The authors thank Dr Ariberto Fassati (UCL, London, UK) for the critical reading of the manuscript and Dr Nicolas Cermakian (Douglas Institute; McGill University) for valuable discussions the project. Finally, the authors acknowledge the key contribution of all study participants for their precious gift of leukapheresis essential for this study.

## Footnote

This work was supported by a grant from the Canadian HIV Cure Enterprise Team Grant (CanCURE 1.0) funded by the Canadian Institutes of Health (CIHR) in partnership with CANFAR and IAS (CanCURE 1.0; # HIG-133050 to PA); the Canadian HIV Cure Enterprise Team Grant (CanCURE 2.0) funded by the CIHR (#HB2-164064); and CIHR project grants to PA (PJT #153052; PJT 178127). Core facilities and PLWH cohorts were supported by the *Fondation du CHUM* and the FRQ-S/AIDS and Infectious Diseases Network. The funding institutions played no role in the design, collection, analysis, and interpretation of data. TRWS received doctoral fellowships from the Université de Montréal and Fonds de Recherche Québec Santé (FRQ-S). JPR holds the Louis Lowenstein Chair in Hematology and Oncology, McGill University.

## Authors’ contributions

DC performed the majority of the experiments, prepared the figures, and wrote the manuscript. YZ, TRWS, and CDNY optimized assays, performed experiments, and prepared figures. HC and YS performed preliminary experiments in support to the study. JPG performed the analysis of RNA-Sequencing data. BB provided protocols for the T7 assay and contributed to study design. JPR provided access to biological samples and clinical information. PA designed the study, analyzed results, prepared figures, and wrote the manuscript. All authors revised the manuscript and agreed with its submission.

## References

1. Davenport MP, Khoury DS, Cromer D, Lewin SR, Kelleher AD, and Kent SJ. Functional cure of HIV: the scale of the challenge. Nat Rev Immunol. 2019;19(1):45–54.

2. Deeks SG, Archin N, Cannon P, Collins S, Jones RB, de Jong M, et al. Research priorities for an HIV cure: International AIDS Society Global Scientific Strategy 2021. Nat Med. 2021;27(12):2085–98.

3. Cohn LB, Chomont N, and Deeks SG. The Biology of the HIV-1 Latent Reservoir and Implications for Cure Strategies. Cell Host Microbe. 2020;27(4):519–30.

4. Sengupta S, and Siliciano RF. Targeting the Latent Reservoir for HIV-1. Immunity. 2018;48(5):872–95.

5. Siliciano JD, and Siliciano RF. Nonsuppressible HIV-1 viremia: a reflection of how the reservoir persists. J Clin Invest. 2020;130(11):5665–7.

6. Siliciano JD, and Siliciano RF. In Vivo Dynamics of the Latent Reservoir for HIV-1: New Insights and Implications for Cure. Annu Rev Pathol. 2022;17:271–94.

7. Mediouni S, Lyu S, Schader SM, and Valente ST. Forging a Functional Cure for HIV: Transcription Regulators and Inhibitors. Viruses. 2022;14(9).

8. Wacleche VS, Landay A, Routy JP, and Ancuta P. The Th17 Lineage: From Barrier Surfaces Homeostasis to Autoimmunity, Cancer, and HIV-1 Pathogenesis. Viruses. 2017;9(10).

9. Planas D, Routy JP, and Ancuta P. New Th17-specific therapeutic strategies for HIV remission. Curr Opin HIV AIDS. 2019;14(2):85–92.

10. Butterfield TR, Landay AL, and Anzinger JJ. Dysfunctional Immunometabolism in HIV Infection: Contributing Factors and Implications for Age-Related Comorbid Diseases. Curr HIV/AIDS Rep. 2020;17(2):125–37.

11. Wagle A, Goerlich E, Post WS, Woldu B, Wu KC, and Hays AG. HIV and Global Cardiovascular Health. Curr Cardiol Rep. 2022.

12. Busman-Sahay K, Starke CE, Nekorchuk MD, and Estes JD. Eliminating HIV reservoirs for a cure: the issue is in the tissue. Curr Opin HIV AIDS. 2021;16(4):200–8.

13. Fert A, Raymond Marchand L, Wiche Salinas TR, and Ancuta P. Targeting Th17 cells in HIV-1 remission/cure interventions. Trends Immunol. 2022;43(7):580–94.

14. Brenchley JM, Paiardini M, Knox KS, Asher AI, Cervasi B, Asher TE, et al. Differential Th17 CD4 T-cell depletion in pathogenic and nonpathogenic lentiviral infections. Blood. 2008;112(7):2826–35.

15. Brenchley JM, and Paiardini M. Immunodeficiency lentiviral infections in natural and non-natural hosts. Blood. 2011;118(4):847–54.

16. Favre D, Lederer S, Kanwar B, Ma ZM, Proll S, Kasakow Z, et al. Critical loss of the balance between Th17 and T regulatory cell populations in pathogenic SIV infection. PLoS Pathog. 2009;5(2):e1000295.

17. Ciccone EJ, Greenwald JH, Lee PI, Biancotto A, Read SW, Yao MA, et al. CD4+ T cells, including Th17 and cycling subsets, are intact in the gut mucosa of HIV-1-infected long-term nonprogressors. J Virol. 2011;85(12):5880–8.

18. Caetano DG, de Paula HHS, Bello G, Hoagland B, Villela LM, Grinsztejn B, et al. HIV-1 elite controllers present a high frequency of activated regulatory T and Th17 cells. PLoS One. 2020;15(2):e0228745.

19. Ivanov, II, McKenzie BS, Zhou L, Tadokoro CE, Lepelley A, Lafaille JJ, et al. The orphan nuclear receptor RORgammat directs the differentiation program of proinflammatory IL-17+ T helper cells. Cell. 2006;126(6):1121–33.

20. Unutmaz D. RORC2: the master of human Th17 cell programming. Eur J Immunol. 2009;39(6):1452–5.

21. Solt LA, Kumar N, Nuhant P, Wang Y, Lauer JL, Liu J, et al. Suppression of TH17 differentiation and autoimmunity by a synthetic ROR ligand. Nature. 2011;472(7344):491–4.

22. Xiao S, Yosef N, Yang J, Wang Y, Zhou L, Zhu C, et al. Small-molecule RORgammat antagonists inhibit T helper 17 cell transcriptional network by divergent mechanisms. Immunity. 2014;40(4):477–89.

23. Withers DR, Hepworth MR, Wang X, Mackley EC, Halford EE, Dutton EE, et al. Transient inhibition of ROR-gammat therapeutically limits intestinal inflammation by reducing TH17 cells and preserving group 3 innate lymphoid cells. Nat Med. 2016;22(3):319–23.

24. Wiche Salinas TR, Zhang Y, Sarnello D, Zhyvoloup A, Marchand LR, Fert A, et al. Th17 cell master transcription factor RORC2 regulates HIV-1 gene expression and viral outgrowth. Proc Natl Acad Sci U S A. 2021;118(48).

25. Lee Y, Awasthi A, Yosef N, Quintana FJ, Xiao S, Peters A, et al. Induction and molecular signature of pathogenic TH17 cells. Nat Immunol. 2012;13(10):991–9.

26. Schnell A, Huang L, Singer M, Singaraju A, Barilla RM, Regan BML, et al. Stem-like intestinal Th17 cells give rise to pathogenic effector T cells during autoimmunity. Cell. 2021;184(26):6281–98 e23.

27. Ramesh R, Kozhaya L, McKevitt K, Djuretic IM, Carlson TJ, Quintero MA, et al. Pro-inflammatory human Th17 cells selectively express P-glycoprotein and are refractory to glucocorticoids. J Exp Med. 2014;211(1):89–104.

28. Burbach KM, Poland A, and Bradfield CA. Cloning of the Ah-receptor cDNA reveals a distinctive ligand-activated transcription factor. Proc Natl Acad Sci U S A. 1992;89(17):8185–9.

29. Fukunaga BN, Probst MR, Reisz-Porszasz S, and Hankinson O. Identification of functional domains of the aryl hydrocarbon receptor. J Biol Chem. 1995;270(49):29270–8.

30. Hooper LV. You AhR what you eat: linking diet and immunity. Cell. 2011;147(3):489–91.

31. Lamas B, Natividad JM, and Sokol H. Aryl hydrocarbon receptor and intestinal immunity. Mucosal Immunol. 2018;11(4):1024–38.

32. Stockinger B, Shah K, and Wincent E. AHR in the intestinal microenvironment: safeguarding barrier function. Nat Rev Gastroenterol Hepatol. 2021;18(8):559–70.

33. Gutierrez-Vazquez C, and Quintana FJ. Regulation of the Immune Response by the Aryl Hydrocarbon Receptor. Immunity. 2018;48(1):19–33.

34. Veldhoen M, Hirota K, Westendorf AM, Buer J, Dumoutier L, Renauld JC, et al. The aryl hydrocarbon receptor links TH17-cell-mediated autoimmunity to environmental toxins. Nature. 2008;453(7191):106–9.

35. Quintana FJ, Basso AS, Iglesias AH, Korn T, Farez MF, Bettelli E, et al. Control of T(reg) and T(H)17 cell differentiation by the aryl hydrocarbon receptor. Nature. 2008.

36. Kimura A, Naka T, Nohara K, Fujii-Kuriyama Y, and Kishimoto T. Aryl hydrocarbon receptor regulates Stat1 activation and participates in the development of Th17 cells. Proc Natl Acad Sci U S A. 2008;105(28):9721–6.

37. Veldhoen M, Hirota K, Christensen J, O’Garra A, and Stockinger B. Natural agonists for aryl hydrocarbon receptor in culture medium are essential for optimal differentiation of Th17 T cells. J Exp Med. 2009;206(1):43–9.

38. Gandhi R, Kumar D, Burns EJ, Nadeau M, Dake B, Laroni A, et al. Activation of the aryl hydrocarbon receptor induces human type 1 regulatory T cell-like and Foxp3(+) regulatory T cells. Nat Immunol. 2010;11(9):846–53.

39. Apetoh L, Quintana FJ, Pot C, Joller N, Xiao S, Kumar D, et al. The aryl hydrocarbon receptor interacts with c-Maf to promote the differentiation of type 1 regulatory T cells induced by IL-27. Nat Immunol. 2010;11(9):854–61.

40. Trifari S, Kaplan CD, Tran EH, Crellin NK, and Spits H. Identification of a human helper T cell population that has abundant production of interleukin 22 and is distinct from T(H)-17, T(H)1 and T(H)2 cells. Nat Immunol. 2009;10(8):864-71.

41. Gagliani N, Vesely MC, Iseppon A, Brockmann L, Xu H, Palm NW, et al. Th17 cells transdifferentiate into regulatory T cells during resolution of inflammation. Nature. 2015;523(7559):221–5.

42. Monteiro P, Gilot D, Le Ferrec E, Lecureur V, N’Diaye M, Le Vee M, et al. AhR- and c-maf-dependent induction of beta7-integrin expression in human macrophages in response to environmental polycyclic aromatic hydrocarbons. Biochem Biophys Res Commun. 2007;358(2):442–8.

43. Kueck T, Cassella E, Holler J, Kim B, and Bieniasz PD. The aryl hydrocarbon receptor and interferon gamma generate antiviral states via transcriptional repression. eLife. 2018;7.

44. Murray IA, Patterson AD, and Perdew GH. Aryl hydrocarbon receptor ligands in cancer: friend and foe. Nature reviews Cancer. 2014;14(12):801–14.

45. Dean JW, and Zhou L. Cell-intrinsic view of the aryl hydrocarbon receptor in tumor immunity. Trends Immunol. 2022;43(3):245–58.

46. Hultquist JF, Schumann K, Woo JM, Manganaro L, McGregor MJ, Doudna J, et al. A Cas9 Ribonucleoprotein Platform for Functional Genetic Studies of HIV-Host Interactions in Primary Human T Cells. Cell reports. 2016;17(5):1438–52.

47. Parrish NF, Gao F, Li H, Giorgi EE, Barbian HJ, Parrish EH, et al. Phenotypic properties of transmitted founder HIV-1. Proc Natl Acad Sci U S A. 2013;110(17):6626–33.

48. Denison MS, and Nagy SR. Activation of the aryl hydrocarbon receptor by structurally diverse exogenous and endogenous chemicals. Annu Rev Pharmacol Toxicol. 2003;43:309–34.

49. Rannug A, and Rannug U. The tryptophan derivative 6-formylindolo[3,2-b]carbazole, FICZ, a dynamic mediator of endogenous aryl hydrocarbon receptor signaling, balances cell growth and differentiation. Crit Rev Toxicol. 2018;48(7):555–74.

50. Zhao B, Degroot DE, Hayashi A, He G, and Denison MS. CH223191 is a ligand-selective antagonist of the Ah (Dioxin) receptor. Toxicol Sci. 2010;117(2):393–403.

51. Macedo AB, Novis CL, De Assis CM, Sorensen ES, Moszczynski P, Huang SH, et al. Dual TLR2 and TLR7 agonists as HIV latency-reversing agents. JCI Insight. 2018;3(19).

52. Kedzierska K, and Crowe SM. Cytokines and HIV-1: interactions and clinical implications. Antiviral chemistry & chemotherapy. 2001;12(3):133–50.

53. Harper J, Ribeiro SP, Chan CN, Aid M, Deleage C, Micci L, et al. Interleukin-10 contributes to reservoir establishment and persistence in SIV-infected macaques treated with antiretroviral therapy. J Clin Invest. 2022;132(8).

54. Wan Q, Kozhaya L, ElHed A, Ramesh R, Carlson TJ, Djuretic IM, et al. Cytokine signals through PI-3 kinase pathway modulate Th17 cytokine production by CCR6+ human memory T cells. J Exp Med. 2011;208(9):1875–87.

55. Finkelshtein D, Werman A, Novick D, Barak S, and Rubinstein M. LDL receptor and its family members serve as the cellular receptors for vesicular stomatitis virus. Proc Natl Acad Sci U S A. 2013;110(18):7306–11.

56. Abdel-Mohsen M, Richman D, Siliciano RF, Nussenzweig MC, Howell BJ, Martinez-Picado J, et al. Recommendations for measuring HIV reservoir size in cure-directed clinical trials. Nat Med. 2020;26(9):1339–50.

57. Zhang Y, Planas D, Raymond Marchand L, Massanella M, Chen H, Wacleche VS, et al. Improving HIV Outgrowth by Optimizing Cell-Culture Conditions and Supplementing With all-trans Retinoic Acid. Frontiers in microbiology. 2020;11:902.

58. Sie C, Kant R, Peter C, Muschaweckh A, Pfaller M, Nirschl L, et al. IL-24 intrinsically regulates Th17 cell pathogenicity in mice. J Exp Med. 2022;219(8).

59. Chen WY, Wang DH, Yen RC, Luo J, Gu W, and Baylin SB. Tumor suppressor HIC1 directly regulates SIRT1 to modulate p53-dependent DNA-damage responses. Cell. 2005;123(3):437–48.

60. Le Douce V, Forouzanfar F, Eilebrecht S, Van Driessche B, Ait-Ammar A, Verdikt R, et al. HIC1 controls cellular- and HIV-1-gene transcription via interactions with CTIP2 and HMGA1. Sci Rep. 2016;6:34920.

61. Burrows K, Antignano F, Bramhall M, Chenery A, Scheer S, Korinek V, et al. The transcriptional repressor HIC1 regulates intestinal immune homeostasis. Mucosal Immunol. 2017;10(6):1518–28.

62. Crowl JT, Heeg M, Ferry A, Milner JJ, Omilusik KD, Toma C, et al. Tissue-resident memory CD8(+) T cells possess unique transcriptional, epigenetic and functional adaptations to different tissue environments. Nat Immunol. 2022;23(7):1121–31.

63. Hsiao F, Frouard J, Gramatica A, Xie G, Telwatte S, Lee GQ, et al. Tissue memory CD4+ T cells expressing IL-7 receptor-alpha (CD127) preferentially support latent HIV-1 infection. PLoS Pathog. 2020;16(4):e1008450.

64. Telwatte S, Lee S, Somsouk M, Hatano H, Baker C, Kaiser P, et al. Gut and blood differ in constitutive blocks to HIV transcription, suggesting tissue-specific differences in the mechanisms that govern HIV latency. PLoS Pathog. 2018;14(11):e1007357.

65. Yukl SA, Kaiser P, Kim P, Telwatte S, Joshi SK, Vu M, et al. HIV latency in isolated patient CD4(+) T cells may be due to blocks in HIV transcriptional elongation, completion, and splicing. Sci Transl Med. 2018;10(430).

66. Zack JA, Kim SG, and Vatakis DN. HIV restriction in quiescent CD4(+) T cells. Retrovirology. 2013;10:37.

67. Descours B, Cribier A, Chable-Bessia C, Ayinde D, Rice G, Crow Y, et al. SAMHD1 restricts HIV-1 reverse transcription in quiescent CD4(+) T-cells. Retrovirology. 2012;9:87.

68. Wiche Salinas TR, Gosselin A, Raymond Marchand L, Moreira Gabriel E, Tastet O, Goulet JP, et al. IL-17A reprograms intestinal epithelial cells to facilitate HIV-1 replication and outgrowth in CD4+ T cells. iScience. 2021;24(11):103225.

69. Wu C, Yosef N, Thalhamer T, Zhu C, Xiao S, Kishi Y, et al. Induction of pathogenic TH17 cells by inducible salt-sensing kinase SGK1. Nature. 2013;496(7446):513–7.

70. Wang R, Campbell S, Amir M, Mosure SA, Bassette MA, Eliason A, et al. Genetic and pharmacological inhibition of the nuclear receptor RORalpha regulates TH17 driven inflammatory disorders. Nat Commun. 2021;12(1):76.

71. Monteiro P, Gosselin A, Wacleche VS, El-Far M, Said EA, Kared H, et al. Memory CCR6+CD4+ T cells are preferential targets for productive HIV type 1 infection regardless of their expression of integrin beta7. J Immunol. 2011;186(8):4618–30.

72. Cicala C, Martinelli E, McNally JP, Goode DJ, Gopaul R, Hiatt J, et al. The integrin alpha4beta7 forms a complex with cell-surface CD4 and defines a T-cell subset that is highly susceptible to infection by HIV-1. Proc Natl Acad Sci U S A. 2009;106(49):20877–82.

73. Nawaz F, Goes LR, Ray JC, Olowojesiku R, Sajani A, Ansari AA, et al. MAdCAM costimulation through Integrin-alpha4beta7 promotes HIV replication. Mucosal Immunol. 2018;11(5):1342–51.

74. Swaminathan G, Nguyen LP, Namkoong H, Pan J, Haileselassie Y, Patel A, et al. The aryl hydrocarbon receptor regulates expression of mucosal trafficking receptor GPR15. Mucosal Immunol. 2021;14(4):852–61.

75. Benechet AP, Menon M, Xu D, Samji T, Maher L, Murooka TT, et al. T cell-intrinsic S1PR1 regulates endogenous effector T-cell egress dynamics from lymph nodes during infection. Proc Natl Acad Sci U S A. 2016;113(8):2182–7.

76. Ancuta P, Autissier P, Wurcel A, Zaman T, Stone D, and Gabuzda D. CD16+ Monocyte-Derived Macrophages Activate Resting T Cells for HIV Infection by Producing CCR3 and CCR4 Ligands. J Immunol. 2006;176(10):5760–71.

77. Jenabian MA, El-Far M, Vyboh K, Kema I, Costiniuk CT, Thomas R, et al. Immunosuppressive Tryptophan Catabolism and Gut Mucosal Dysfunction Following Early HIV Infection. J Infect Dis. 2015;212(3):355–66.

78. Chen J, Xun J, Yang J, Ji Y, Liu L, Qi T, et al. Plasma Indoleamine 2,3-Dioxygenase Activity Is Associated With the Size of the Human Immunodeficiency Virus Reservoir in Patients Receiving Antiretroviral Therapy. Clin Infect Dis. 2019;68(8):1274–81.

79. Dai L, Lidie KB, Chen Q, Adelsberger JW, Zheng X, Huang D, et al. IL-27 inhibits HIV-1 infection in human macrophages by down-regulating host factor SPTBN1 during monocyte to macrophage differentiation. J Exp Med. 2013;210(3):517–34.

80. Lederman MM, Funderburg NT, Sekaly RP, Klatt NR, and Hunt PW. Residual immune dysregulation syndrome in treated HIV infection. Advances in immunology. 2013;119:51–83.

81. Sun X, Qu Q, Lao Y, Zhang M, Yin X, Zhu H, et al. Tumor suppressor HIC1 is synergistically compromised by cancer-associated fibroblasts and tumor cells through the IL-6/pSTAT3 axis in breast cancer. BMC Cancer. 2019;19(1):1180.

82. Pagans S, Pedal A, North BJ, Kaehlcke K, Marshall BL, Dorr A, et al. SIRT1 regulates HIV transcription via Tat deacetylation. PLoS Biol. 2005;3(2):e41.

83. Pinzone MR, Cacopardo B, Condorelli F, Di Rosa M, and Nunnari G. Sirtuin-1 and HIV-1: an overview. Curr Drug Targets. 2013;14(6):648–52.

84. Limagne E, Thibaudin M, Euvrard R, Berger H, Chalons P, Vegan F, et al. Sirtuin-1 Activation Controls Tumor Growth by Impeding Th17 Differentiation via STAT3 Deacetylation. Cell reports. 2017;19(4):746–59.

85. Mameli G, Deshmane SL, Ghafouri M, Cui J, Simbiri K, Khalili K, et al. C/EBPbeta regulates human immunodeficiency virus 1 gene expression through its association with cdk9. J Gen Virol. 2007;88(Pt 2):631–40.

86. Wagner TA, McLaughlin S, Garg K, Cheung CY, Larsen BB, Styrchak S, et al. HIV latency. Proliferation of cells with HIV integrated into cancer genes contributes to persistent infection. Science. 2014;345(6196):570–3.

87. Maldarelli F, Wu X, Su L, Simonetti FR, Shao W, Hill S, et al. HIV latency. Specific HIV integration sites are linked to clonal expansion and persistence of infected cells. Science. 2014;345(6193):179–83.

88. Cesana D, Santoni de Sio FR, Rudilosso L, Gallina P, Calabria A, Beretta S, et al. HIV-1-mediated insertional activation of STAT5B and BACH2 trigger viral reservoir in T regulatory cells. Nat Commun. 2017;8(1):498.

89. Zhou YH, Sun L, Chen J, Sun WW, Ma L, Han Y, et al. Tryptophan Metabolism Activates Aryl Hydrocarbon Receptor-Mediated Pathway To Promote HIV-1 Infection and Reactivation. mBio. 2019;10(6).

90. Ohtake F, Baba A, Takada I, Okada M, Iwasaki K, Miki H, et al. Dioxin receptor is a ligand-dependent E3 ubiquitin ligase. Nature. 2007;446(7135):562–6.

91. Das B, Dobrowolski C, Luttge B, Valadkhan S, Chomont N, Johnston R, et al. Estrogen receptor-1 is a key regulator of HIV-1 latency that imparts gender-specific restrictions on the latent reservoir. Proc Natl Acad Sci U S A. 2018;115(33):E7795–E804.

92. Luecke-Johansson S, Gralla M, Rundqvist H, Ho JC, Johnson RS, Gradin K, et al. A Molecular Mechanism To Switch the Aryl Hydrocarbon Receptor from a Transcription Factor to an E3 Ubiquitin Ligase. Mol Cell Biol. 2017;37(13).

93. Gosselin A, Wiche Salinas TR, Planas D, Wacleche VS, Zhang Y, Fromentin R, et al. HIV persists in CCR6+CD4+ T cells from colon and blood during antiretroviral therapy. AIDS. 2017;31(1):35–48.

94. Planas D, Fert A, Zhang Y, Goulet JP, Richard J, Finzi A, et al. Pharmacological Inhibition of PPARy Boosts HIV Reactivation and Th17 Effector Functions, While Preventing Progeny Virion Release and de novo Infection. Pathogens & immunity. 2020;5(1):177–239.

95. Ochsenbauer C, Edmonds TG, Ding H, Keele BF, Decker J, Salazar MG, et al. Generation of transmitted/founder HIV-1 infectious molecular clones and characterization of their replication capacity in CD4 T lymphocytes and monocyte-derived macrophages. J Virol. 2012;86(5):2715–28.

96. Planas D, Zhang Y, Monteiro P, Goulet JP, Gosselin A, Grandvaux N, et al. HIV-1 selectively targets gut-homing CCR6+CD4+ T cells via mTOR-dependent mechanisms. JCI Insight. 2017;2(15).

97. Grajkowska LT, Ceribelli M, Lau CM, Warren ME, Tiniakou I, Nakandakari Higa S, et al. Isoform-Specific Expression and Feedback Regulation of E Protein TCF4 Control Dendritic Cell Lineage Specification. Immunity. 2017;46(1):65–77.

98. Cleret-Buhot A, Zhang Y, Planas D, Goulet JP, Monteiro P, Gosselin A, et al. Identification of novel HIV-1 dependency factors in primary CCR4(+)CCR6(+)Th17 cells via a genome-wide transcriptional approach. Retrovirology. 2015;12:102.

