## Supplemental Figures 1-12 for "Identification of Aryl Hydrocarbon Receptor as a Barrier to HIV-1 Infection and Outgrowth in CD4^+^ T-Cells"

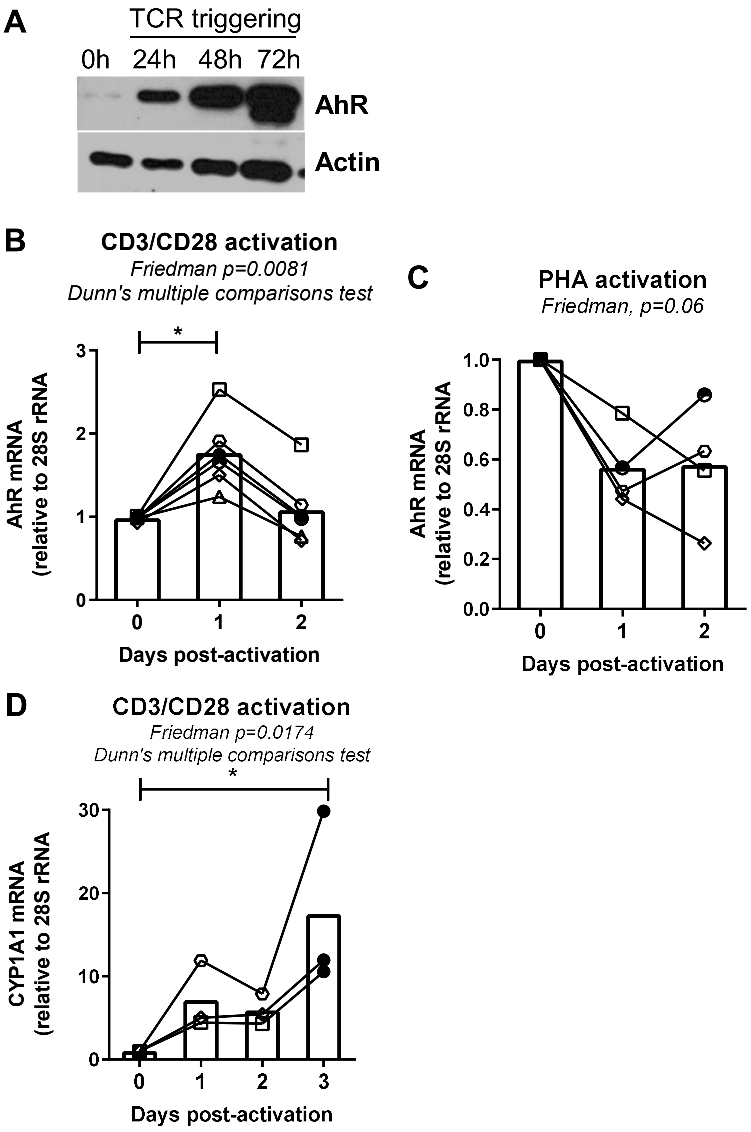


**Supplemental Figure 1: AhR expression and activation is induced by TCR triggering but not PHA stimulation in memory CD4⁺ T-cells.** Memory CD4⁺ T-cells isolated from PBMCs of HIV-uninfected individuals were stimulated with CD3/CD28 Abs for 1-3 days or PHA for 2 days. AhR protein expression was measured by western blotting *ex vivo* and at days 1, 2, and 3 post-TCR triggering **(A)**. In parallel, total RNA was extracted from cells activated with CD3/CD28 Abs for 1-3 days (**B)** or PHA (10 µg/ml) for 2 days (**C)** and used for the quantification of AhR mRNA **(B-C)** and CyP1A1 mRNA **(D)** by real-time RT-PCR (n=4-6). Friedman test p-values, with Dunn's multiple comparisons indicated on the graphs.

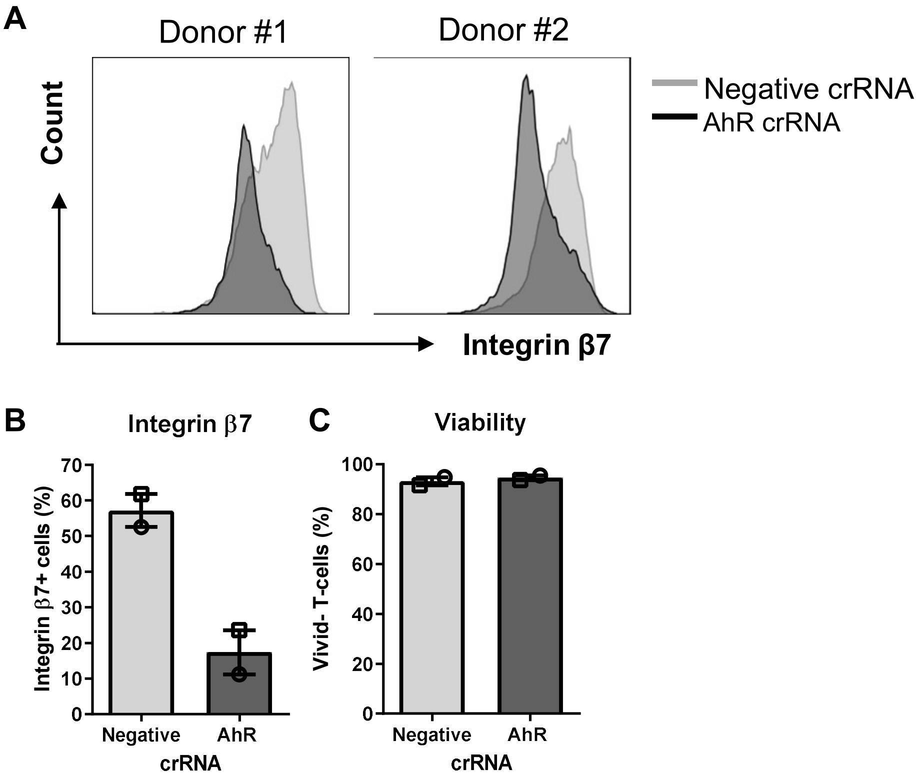


**Supplemental Figure 2: CRISPR/Cas9-mediated AhR KO downregulates integrin β7 and IL-22 expression in memory CD4⁺ T-cells.** Memory CD4⁺ T-cells isolated from PBMCs of HIV-uninfected individuals were stimulated with CD3/CD28 Abs and nucleofected either with negative crRNA or with AhR crRNA, as in Figure 1. After four days, cells were analyzed by flow cytometry for the surface expression of integrin β7 (ITGB7) (A-B) and cell viability using the viability dye aqua vivid (C). Shown are results from two different HIV-negative individuals, with each symbol representing a different donor.

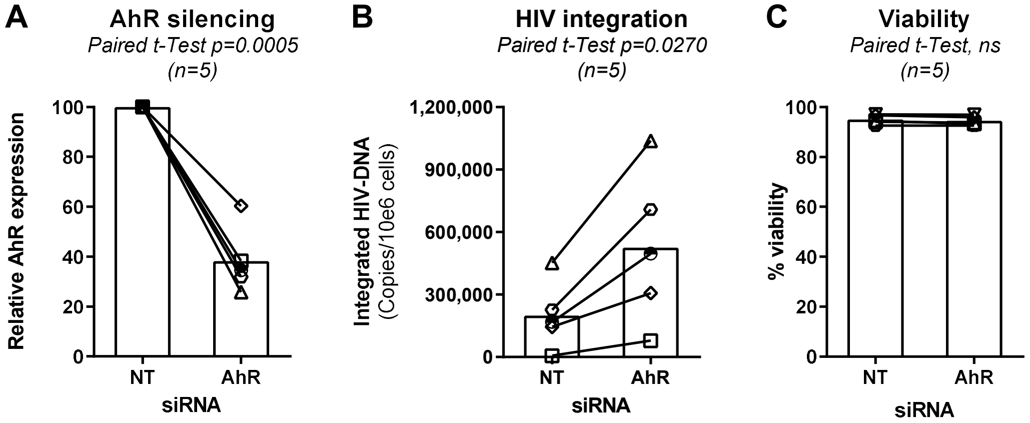


**Supplemental Figure 3: Small interfering RNA against AhR increases HIV-DNA integration in memory $CD4^+$ T-cells.** Memory $CD4^+$ T-cells isolated from the PBMCs of HIV-uninfected individuals were stimulated *via* CD3/CD28 for 3 days. Then, cells were nucleofected with non-targeting (NT) and AhR-targetting small interfering RNA (siRNA). Nucleofected cells were cultured in the presence of IL-2 (5 ng/ml) for 24 hours and then exposed to replication-competent NL4.3BaL HIV-1 strain for 3 hours. Unbound virus was excluded by extensive washing. Cells were then cultured in the presence of IL-2 for 3 other days. Shown are AhR mRNA expression as measured by real-time RT-PCR **(A)**; levels of integrated HIV-DNA quantified by nested real-time PCR in cells **(B)**; and cell viability measured using the viability dye **(C)** upon siRNA silencing. Results were generated with cells from n=5 different HIV-uninfected individuals. Each symbol represents one different donor. Paired t-test p-values are indicated on the graphs.

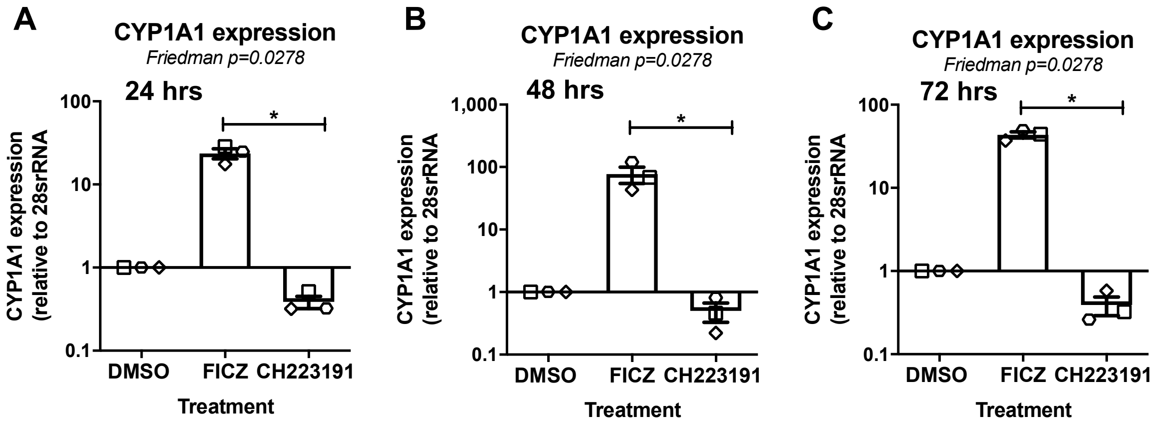


**Supplemental Figure 4: Pharmacological targeting of AhR pathway modulates CYP1A1 mRNA expression.** Memory CD4$^+$ T-cells from the PBMCs of HIV-uninfected individuals were stimulated with CD3/CD28 Abs in the presence or the absence of FICZ (100 nM) or CH223191 (10 μM) for 24 (A), 48 (B) and 72 hours (C). CYP1A1 mRNA expression was measured by RT-PCR, with 28S rRNA used as control housekeeping gene. Shown are CYP1A1 mRNA expression relative to DMSO levels (DMSO considered 1). Results were generated with cells from n=3 different donors. Friedman test p-values, with Dunn's multiple comparisons indicated on the graphs.

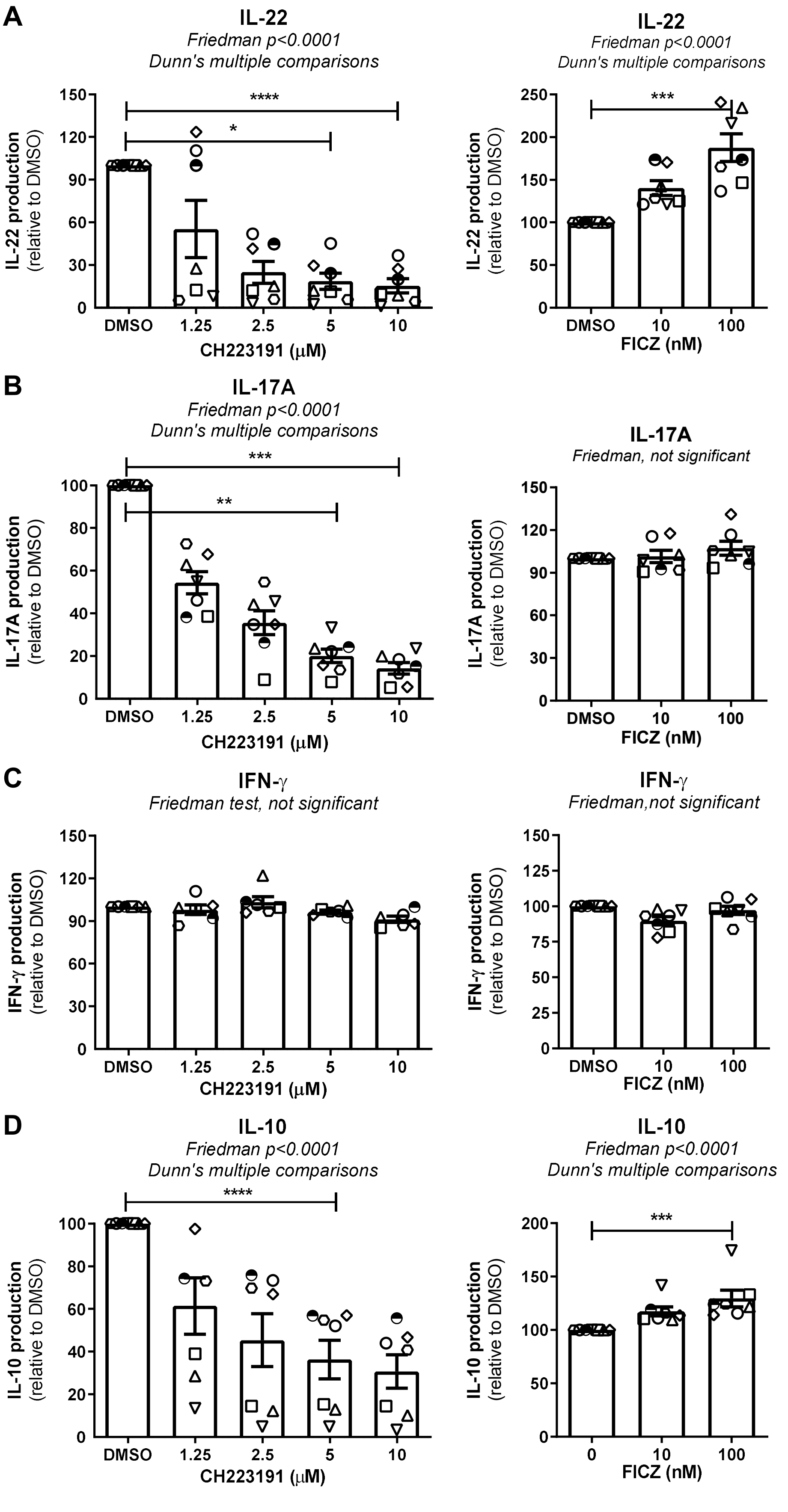


Chatterjee *et al.,* Supplemental Figure 5

**Supplemental Figure 5: Pharmacological targeting of AhR modulates IL-22, IL-17A and IL-10 but not IFN-γ expression in primary CD4⁺ T-cells.** Memory CD4⁺ T-cells isolated as described in Figure 1A were stimulated with CD3/CD28 Abs in the presence or the absence of different concentrations of the AhR antagonist CH223191 (1.25, 2.5, 5 or 10 μM) or agonist FICZ (10 and 100 nM). Shown are levels of IL-22 **(A)**, IL-17A **(B)**, IFN-γ **(C)**, and IL-10 **(D)** measured by ELISA in cell culture supernatants collected from cells exposed to CH223191 (**left panels**) and FICZ (**right panels**) at day 3 post-TCR triggering. Results were generated with cells from n=7 HIV-uninfected individuals. Each symbol represents one donor. Shown are the Friedman test p-values, and the significance of Dunn's multiple comparisons (*, $p < 0.05$; **, $p < 0.01$; ***, $p < 0.001$).

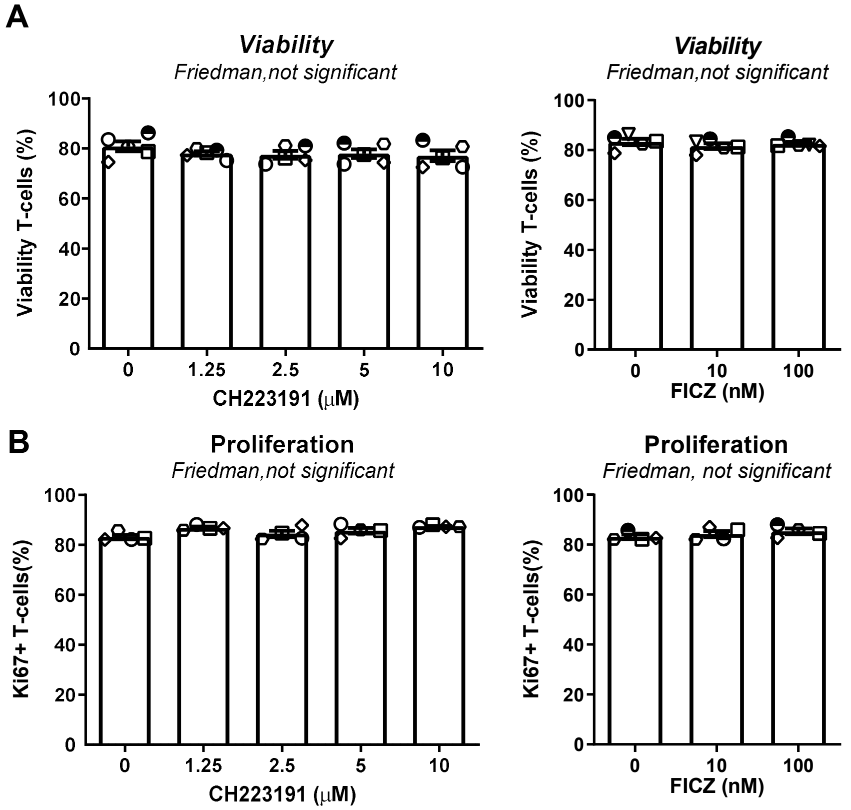


**Supplemental Figure 6: Pharmacological targeting of AhR pathway does not affect cell viability and proliferation**: Memory CD4$^+$ T-cell were isolated from HIV-uninfected individuals and stimulated with CD3/CD28 Abs in the presence or the absence of CH223191 (1.25, 2.5, 5 or 10 μM) or FICZ (10 and100 nM) for 3 days, as in Figure 3. Shown are cell viability was measured by staining using the viability dye aqua vivid **(A)**, as well as cell cycle progression measured by intranuclear staining with Ki-67 Abs (a surrogate marker of cell proliferation) **(B)**. Results were generated with cells from n=4-5 HIV-uninfected individuals. Friedman test p-values, with Dunn's multiple comparisons indicated on the graphs.

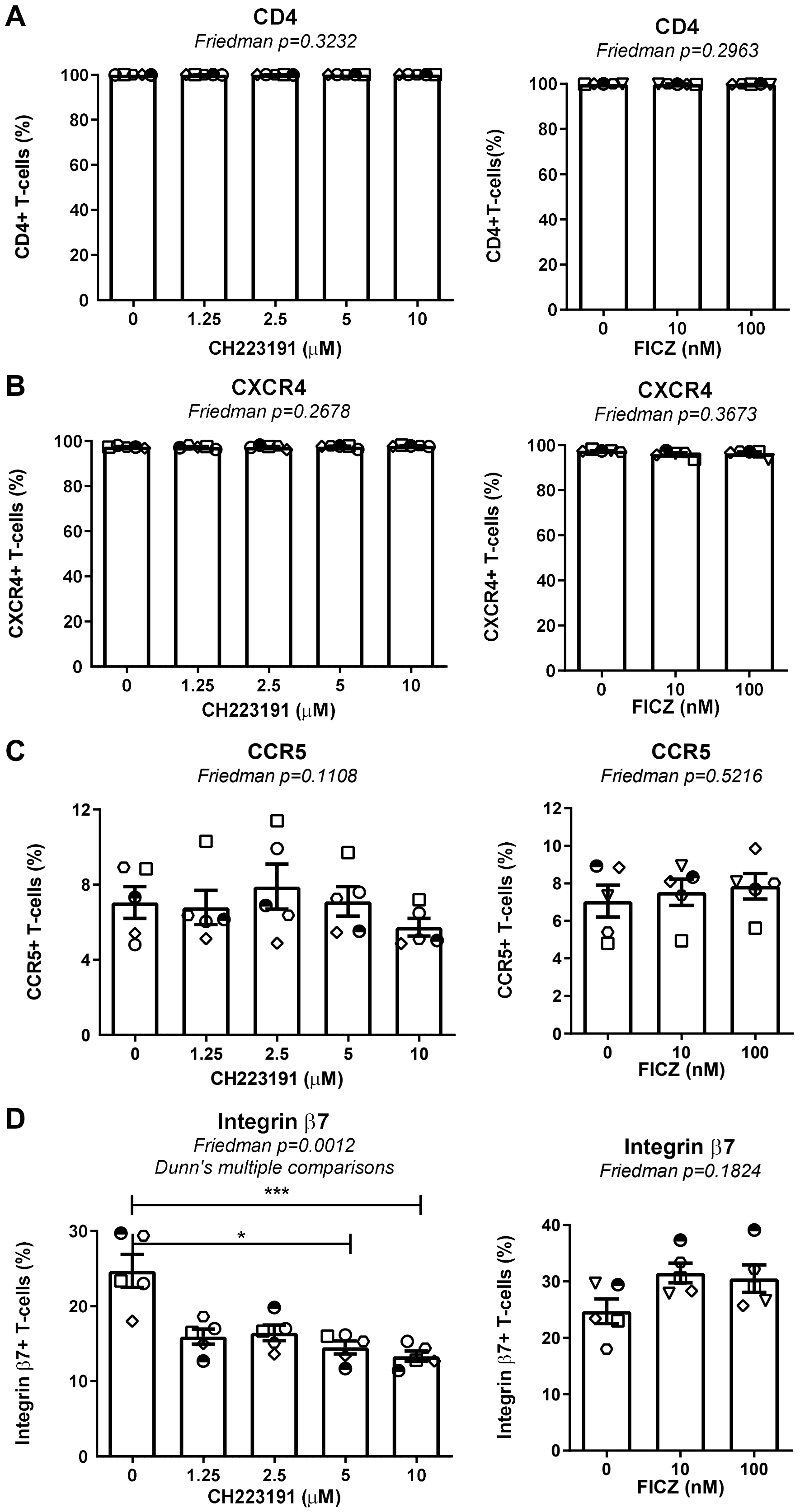


Chatterjee *et al.,* Supplemental Figure 7

**Supplemental Figure 7: AhR antagonism downregulates integrin β7 expression without modulating the expression of the HIV-1 receptor CD4 or co-receptors CCR5 and CXCR4**. Memory $CD4^+$ T-cells were activated *via* CD3/CD28 and cultured in the presence or the absence of different concentrations of CH223191 **(left panels)** or FICZ **(right panels)**, as in Figure 3. Cells were then analyzed by flow cytometry for the surface expression of CD4 **(A),** CXCR4 **(B)**, CCR5 **(C),** and integrin β7 **(D).** Results were generated with cells from n=7 HIV-uninfected individuals. Friedman test p-values, with Dunn's multiple comparisons indicated on the graphs.

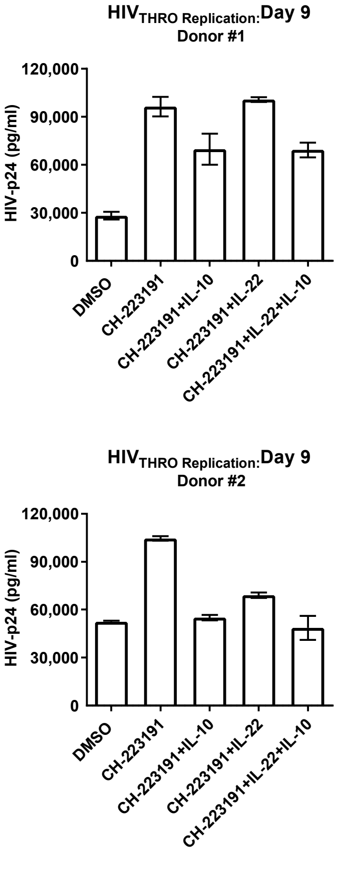


**Supplemental Figure 8: IL-10 and IL-22 supplementation counteracts the proviral effects of CH223191:** Memory CD4$^+$ T-cell were isolated from HIV-uninfected individuals, stimulated with CD3/CD28 Abs and cultured in the presence or the absence of CH223191, as well as human recombinant IL-10 (10 ng/ml) and/or IL-22 (25 ng/ml) for 3 days. Cells were exposed to the replication competent T/F $HIV_{THRO}$ for 3 hours and cultured for additional 9 days. Media containing CH223191 and cytokines was refreshed every 3 days. Shown are HIV-p24 levels measured by ELISA in cell-culture supernatants at day 9 post-infection. Experiments were performed with cells from 2 different donors. Results represent mean±SD of ELISA triplicate values.

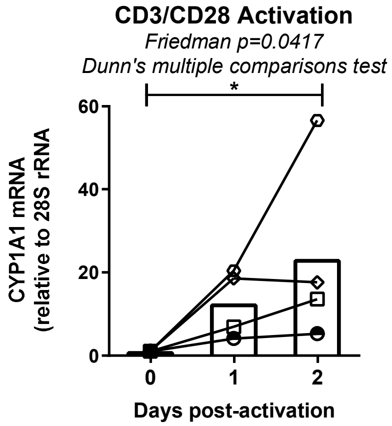


**Supplemental Figure 9: TCR triggering increases CYP1A1 expression in memory CD4⁺ T-cells of ART-treated PLWH.** Memory CD4⁺ T-cell were isolated from ART-treated PLWH and stimulated with CD3/CD28 Abs for up to 2 days. CYP1A1 mRNA expression was measured by RT-PCR and normalized to 28S rRNA levels. Results were generated with cells from n=4 different donors. Friedman test p-values, with Dunn's multiple comparisons indicated on the graphs.

**A**

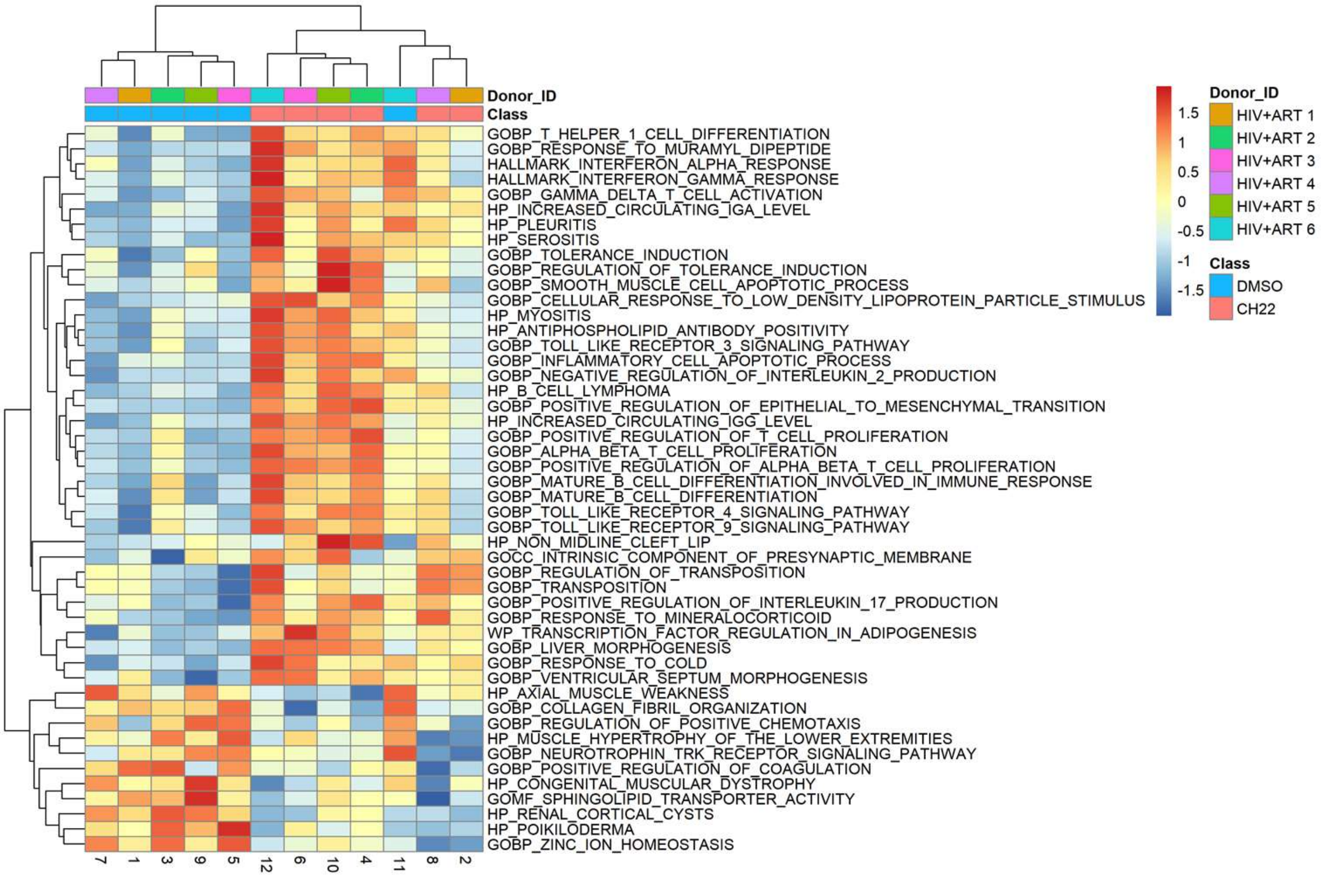


**Supplementary Figure 10: GSVA identifies pathways modulated by CH223191 in memory CD4⁺ T-cells of ART-treated PLWH**: Heatmap representing the gene set variation analysis (GSVA) for all the pathways with a FDR-adjusted p-value <0.05. Heatmap represents pathways included in MSigDB: Heatmap cells are scaled by the expression level z-scores for each probe individually. Results from each donor are indicated with a different color code (n=6). Shown are top regulated pathways **(A)** and individual genes related to selected top regulated pathways **(B).**

**B**

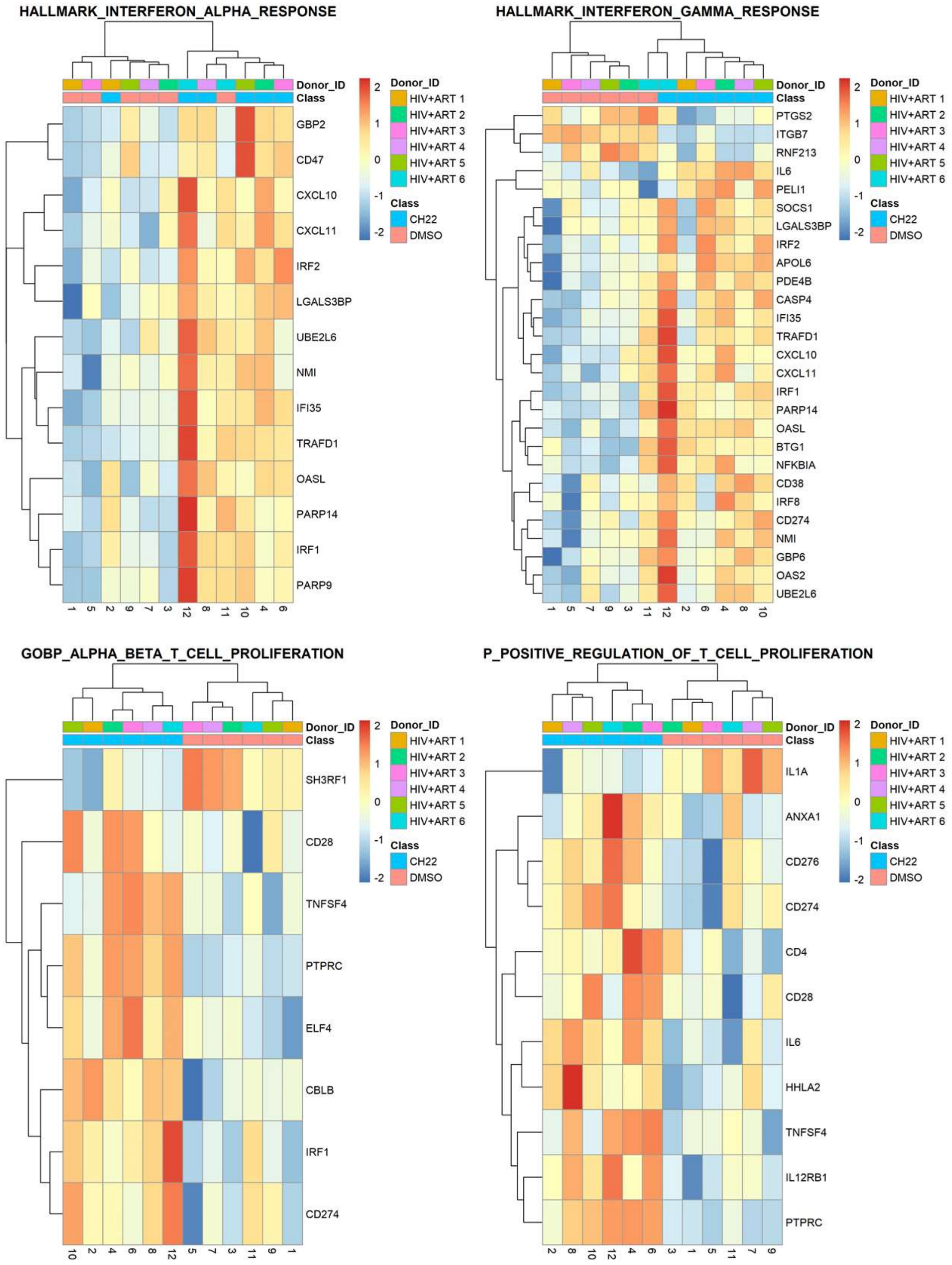


Chatterjee *et al.,* Supplemental Figure 10 - continued

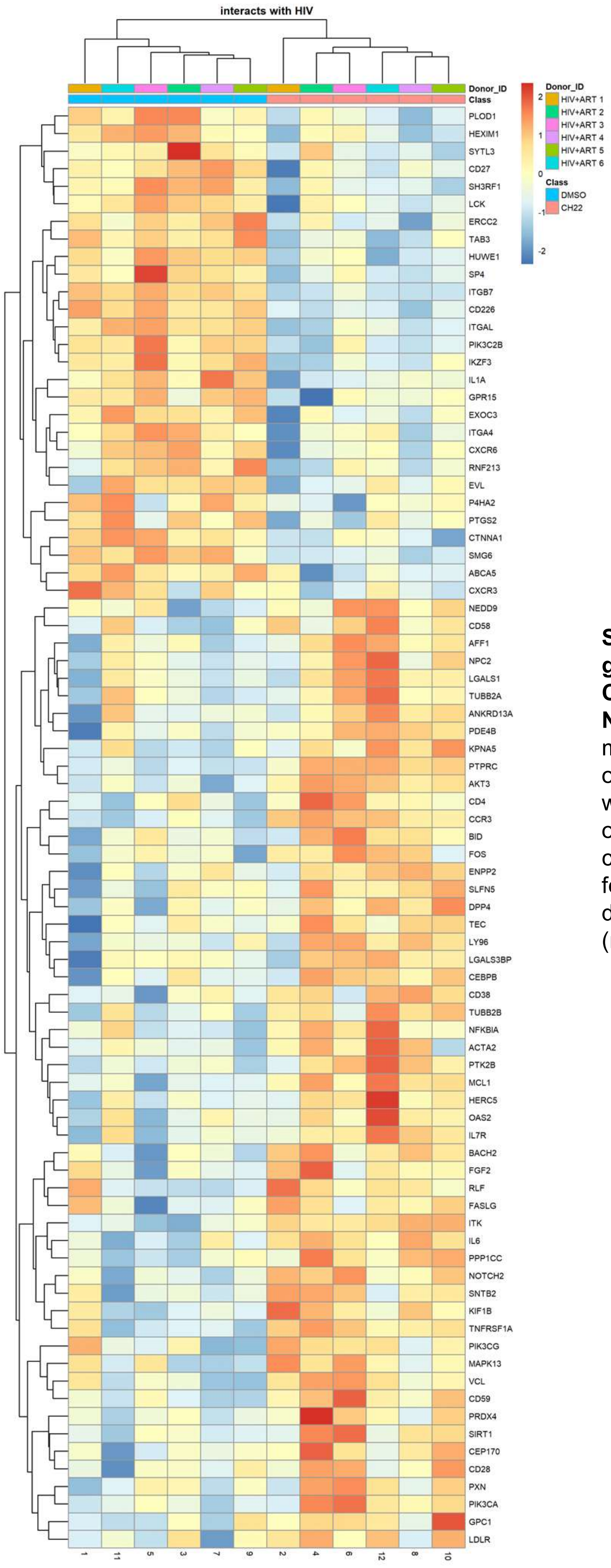


**Supplementary Figure 11: Meta-analysis of genes modulated by CH223191 in memory CD4+ T-cells of ART-treated PLWH using the NCBI HIV interaction database.** Transcripts modulated by CH223191 in memory CD4⁺ T cells of ART-treated PLWH ($p<0.05$; FC cut-off 1.3) were matched to the lists of human genes included on the NCBI HIV interaction database. Heat- map cells are scaled by the expression level z-scores for each probe individually. Results from each donor are indicated with a different color code (n=6).

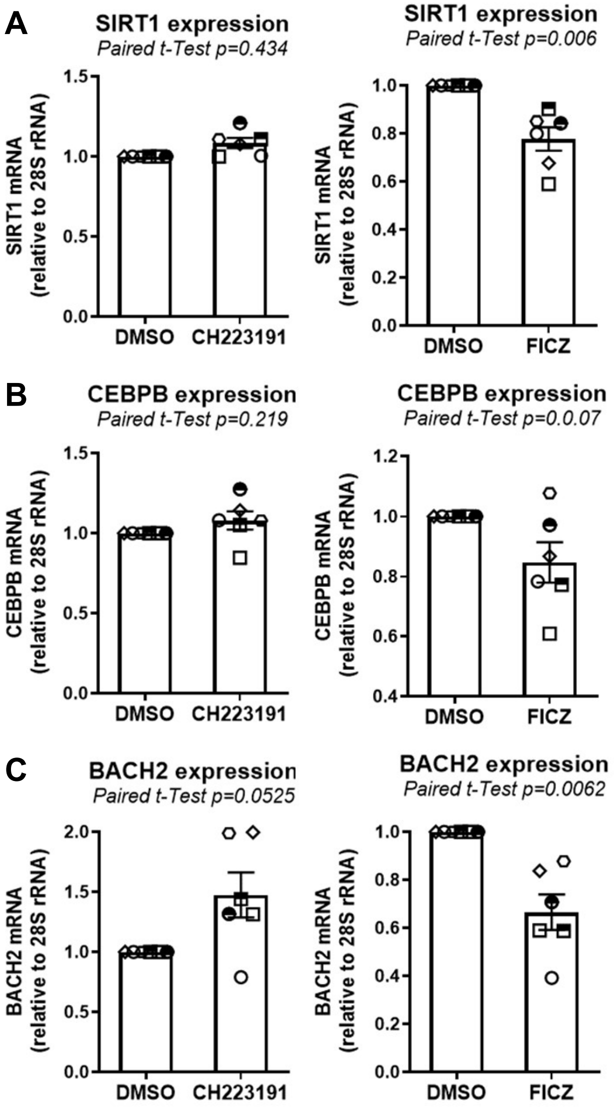


**Supplementary Figure 12: Pharmacological targeting of the AhR pathway modulates expression of SIRT1, CEBPB and BACH2 mRNA:** Briefly, memory $CD4^+$ T cells isolated from the PBMCs of ART-treated PLWH (n=6) were stimulated by CD3/CD28 Abs and cultured in the presence or the absence of CH223191 (10 µM) and FICZ (100 nM) for 18 hours. Total RNA was extracted, and expression levels of SIRT1 **(A)**, CEBPB **(B)** and BACH2 **(C)** transcripts were quantified by qPCR. Each symbol represents one individual donor: bars represent median values. Wilcoxon matched pairs signed rank test are indicated on the graphs.
