## Supplemental Table 1 for "Identification of Aryl Hydrocarbon Receptor as a Barrier to HIV-1 Infection and Outgrowth in CD4^+^ T-Cells"

**Supplemental Table 1**: Clinical parameters of HIV-infected study participants receiving viral suppressive antiretroviral therapy (ART).

| Participant ID | Sex | CD4 counts# | CD8 counts# | Plasma viral load& | Time since infection* | ART | Time of aviremia* |
|---|---|---|---|---|---|---|---|
| **ART #1** | M | 398 | 775 | <40 | 154 | Complera | 22 |
| **ART #2** | M | 542 | 803 | <40 | 13 | Stribild | NA |
| **ART #3** | M | 458 | 899 | <40 | 227 | Truvada/viramune | NA |
| **ART #4** | M | 841 | 1322 | <40 | 149 | Sustiva/ Truvada | NA |
| **ART #5** | M | 598 | 605 | <40 | 80 | Stribild | NA |
| **ART #6** | M | 425 | 1156 | <40 | 182 | Atripla | NA |
| **ART #7** | M | 908 | 854 | <40 | 89 | Stribild | 70 |

#, cells/µl; &, HIV RNA copies per ml plasma; *, months; ART, antiretroviral therapy; NA, information not available; ND, not detected
