## Supplemental Table 2 for "Identification of Aryl Hydrocarbon Receptor as a Barrier to HIV-1 Infection and Outgrowth in CD4^+^ T-Cells"

**Supplemental Table 2: Antibodies used for flow cytometry analysis**

| | Fluorochrome | Clone | Vendor |
|---|---|---|---|
| CD3 | Pacific blue | UCHT1 | BD Pharmingen (San Diego, CA, USA) |
| CD4 | AlexaFluor700 | RPAT4 | |
| CCR6 | PE | 11A9 | |
| CCR5 | PE | 2D7/CCR5 | |
| IFN-γ | AlexaFluor700 | B27 | |
| CXCR4 | PE | 12G5 | |
| Ki67 | FITC | B56 | |
| Integrinβ7 | FITC | FIB504 | eBioscience(San Diego,CA, USA) |
| CD56 | FITC | MEM188 | |
| IL-17A | PE | eBio64BEC17 | |
| IL-22 | PE-Cyanine7 | 22URTI | |
| CD8 | FITC | BW135/80 | Miltenyi Biotech (Auburn, CA, USA) |
| CD19 | FITC | LT19 | |
| CD45RA | APCeFluor780 | HI100 | Invitrogen (Waltham, MA, USA) |
| HIV-p24 | FITC | KC57 | Beckman Coulter (Brea, CA, USA) |
